# Tagging enhances histochemical and biochemical detection of Ran Binding Protein 9 *in vivo* and reveals its interaction with Nucleolin

**DOI:** 10.1101/745174

**Authors:** Shimaa Soliman, Aaron E. Stark, Miranda Gardner, Sean W. Harshman, Chelssie C. Breece, Foued Amari, Arturo Orlacchio, Min Chen, Anna Tessari, Jennifer A. Martin, Rosa Visone, Michael A. Freitas, Krista M.D. La Perle, Dario Palmieri, Vincenzo Coppola

**Author notes:** These authors equally contributed to this work (Shimaa Soliman, Aaron E. Stark). To whom correspondence should be addressed. Vincenzo Coppola, MD Assistant Professor, Cancer Biology & Genetics 988 Biomedical Research Tower, Wexner Medical Center The Ohio State University and Comprehensive Cancer Center 460 West 12th Avenue Columbus, OH 43210.

## Abstract

The lack of tools to reliably detect RanBP9 *in vivo* has significantly hampered progress in understanding the biological functions of this scaffold protein. We report here the generation of a novel mouse strain, RanBP9-TT, in which the endogenous protein is fused with a double (V5-HA) epitope tag at the C-terminus. We show that the double tag does not interfere with the essential functions of RanBP9. In contrast to RanBP9 constitutive knock-out animals, RanBP9-TT mice are viable, fertile and do not show any obvious phenotype. The V5-HA tag allows unequivocal detection of RanBP9 both by IHC and WB. Importantly, immunoprecipitation and mass spectrometry analyses reveal that the tagged protein pulls down known interactors of wild type RanBP9. Thanks to the increased detection power, we are also unveiling a novel interaction with Nucleolin, a prominent nucleolar protein.

In summary, we report the generation of a new mouse line in which RanBP9 expression and interactions can be reliably studied by the use of commercially available αtag antibodies. The use of this line will help to overcome some of the existing limitations in the study of RanBP9 and potentially reveal novel functions of this protein *in vivo* such as those linked to Nucleolin.

## Introduction

RANBP9 is a scaffold protein whose biological functions are not completely understood^1^. Since its initial description, it has been reported to interact with a number of proteins that are located in different cellular compartments and have different biological functions^1^. In many instances RANBP9 expression affects the half life and stability of other proteins ^1^. This function is in accordance with the model in which RanBP9 is part of an E3 ligase multi-subunit structure called the CTLH (C-Terminal to LisH) complex that can induce target proteins degradation via the proteasome pathway. This E3 ligase structure is believed to be the equivalent of the GID (Glucose-Induced degradation Deficient) complex in *S. cerevisiae*, which is involved in responding to changes of nutrient availability in the microenvironment ^2–5^. Although not well studied in mammals, the human CTLH complex has been recently reported to be a heterodecameric structure that, in addition to RANBP9, includes its paralog RANBP10 and nine other poorly studied proteins (ARMC8, GID4, GID8, MAEA, MKLN1, RMND5A, RMND5B, WDR26, and YPEL5) ^5–8^.

Objective difficulties impair the investigation of the cellular pathways and mechanisms in which RANBP9 takes part. Among other issues, the limited specificity and affinity of commercially available antibodies pose significant technical challenges. The existence of a paralog named RANBP10 ^9, 10^ with high similarity in long parts of the protein limits the RANBP9 specific sequence available to raise antibodies. In addition, some of the sequences might not have biochemical features ideal to induce a robust humoral immune response. Therefore, many of the commercially available αRANBP9 antibodies fall short in their detection power, reliability, and/or specificity.

We have previously used the only existing antibody validated by the Human Protein Atlas (HPA050007; www.proteinatlas.org) for immunohistochemical (IHC) detection of human RANBP9 in formalin-fixed paraffin embedded Non-Small Cell Lung Cancer (NSCLC) specimens ^11^. Although specific for human RANBP9, in our hands HPA050007 does not seem to recognize mouse RanBP9 with similar affinity, especially in Western Blot (WB) in which signal-to-noise ratio is low. In addition, WB analysis often shows the presence of additional bands at lower molecular weight that might be due to non-specific binding.

We have used and will continue to use human cell lines for *in vitro* studies. However, to better recapitulate organismal physiology, a significant part of our ongoing studies on RANBP9 involvement in tumor development and response to therapy necessarily takes advantage of *in vivo* murine models. In this regard, we had previously generated the constitutive RanBP9 knockout (KO) animal. On a hybrid C57Bl/6 x S129 genetic background, most homozygous KO mice were dying hours after birth. A small cohort of survivors showed small body size and severe sterility in both males and females ^12^. Using reagents from the International Mouse Phenotyping Consortium (http://www.mousephenotype.org/; IKMC project nr: 44910), we have now engineered the conditional KO mouse that allows the study of RanBP9 loss of function *in vivo*, which recapitulates the phenotype of the constitutive KO when the gene is ubiquitously deleted ^13^. However, to overcome the limitations in the detection of endogenous RanBP9, we have decided to engineer a novel RanBP9 strain in which the wild type protein is fused to commonly used tags against which there are reliable and commercially available antibodies.

Here, we report the generation by CRISPR/Cas9 of the RanBP9-TT (= double Tag = TT) mouse line in which we have inserted a double epitope tag (V5-HA) at the C-terminus of the protein. We show that tagging of RanBP9 allows unequivocal detection of the protein both by IHC and biochemistry without affecting its essential biological functions. Thanks to the superior detection power linked to the use of αtag antibodies we found a previously unknown interaction of RanBP9 with the nucleolar protein Nucleolin. Because we knocked in the tags leaving intact the 3’-untranslated region (UTR) of the RanBP9 genomic locus, the expression of the protein faithfully recapitulates the wild type (WT) expression. Therefore, the RanBP9-TT strain becomes a powerful tool to dissect *in vivo* the biology related to RanBP9 functions allowing its unequivocal detection in murine tissues and cells.

## Results

### Generation of the RanBP9-TT animals

We used CRISPR/Cas9 to knock-in the double tag V5-HA at the C-terminus of RanBP9 (**Figure 1; Figure S1**). For targeting purposes, we employed the online Benchling software (https://www.benchling.com/). We selected the guide RNA (sgRNA) with the best specificity and efficiency scores closest to the insertion site before the stop codon (**Figure 1A and Figure S1A-C**). Pure C57Bl/6Tac WT fertilized eggs were used for the generation of founders (F0) mice. Two F0 animals (#1 and #2) were selected for further breeding and propagation of the RanBP9-TT colony. Both founders produced progeny (F1 mice) positive for the correct insertion of the double tag and animals from both lines were used for this work. Sanger sequencing showed that F1 animals from both founder lines contained the correct in-frame insertion of the V5-HA double tag (**Figure 1C**). In order to mitigate the presence of CRISPR/Cas9 potential off-target effects, we crossed F1 animals a second time to wild type C57Bl/6Tac mice to generated F2 progeny that were used for experimental purposes.

**Figure 1.**
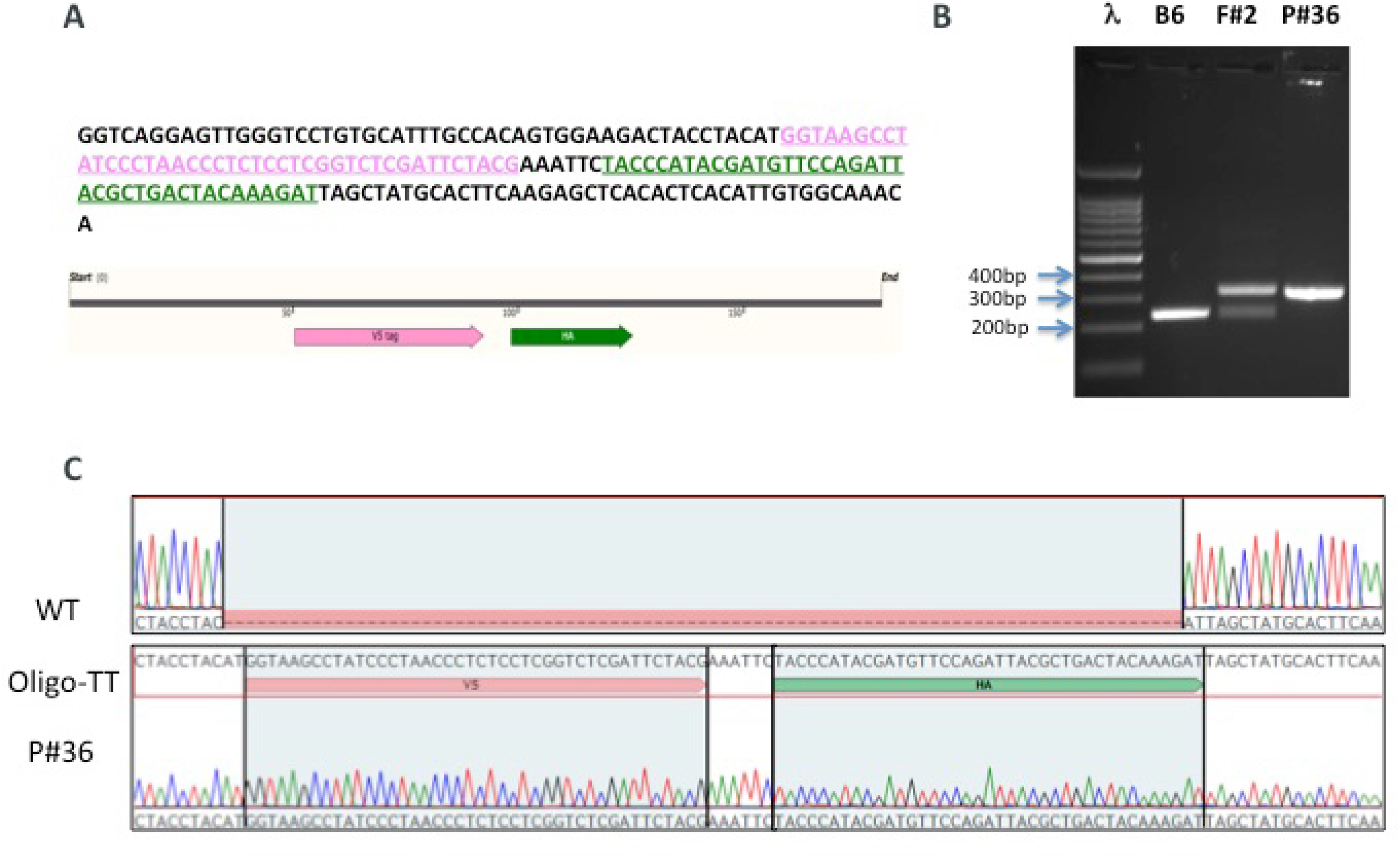
Generation of the RanBP9-TT mouse model by CRISPR/Cas9. **A**) 180bp single strand oligo DNA (ssODN) used as donor to recombine the V5 (<u>PINK</u>) and the HA (<u>GREEN</u>) tags into the C-terminus of RanBP9. **B**) Representative PCR screening results from tail DNA of WT C57Bl/6 (negative control) RanBP9-TT Founder #2 (F#2), and homozygous RanBP9-TT pup number 36 (P#36). Results are congruent with prediction shown in Figure S1D. **C**) Sanger-sequencing results from homozygous pup number 36 compared to C57Bl/6 WT and ssODN shown in A.

These results show that the V5-HA double tag at the C-terminus of endogenous RanBP9 was successfully inserted as designed.

### Addition of tag at the C-terminus does not cause lethality or infertility

On a mixed C57Bl/6 x S129 background using gene-trapped ES cells from the Baygenomics consortium ^14^, homozygous inactivation of RanBP9 causes early postnatal lethality in mice ^12^. The pure C57Bl/6 background seems to worsen the phenotype and homozygous KO animals immediately after birth are rarely found, if any ^15^.

We observed that RanBP9-TT homozygous knock-in (KI) mice are viable and do not show any obvious phenotype. They are born at Mendelian ratio, and there are no differences in the number of females and males born. Adult animals do not show any gross anatomical or histological abnormalities. Most importantly, homozygous RanBP9-TT males and females were bred to each other or to heterozygous and WT C57Bl/6Tac mice. Mice of both genders were able to reproduce in similar numbers to WT controls (**Table 1**). Testes and ovaries of RanBP9-TT mice show histological features similar to WT animals (**Figure 2 and Figures S2, S3, S4**). All together, these results show that the insertion of the V5-HA tag at the C-terminus of RanBP9 does not interfere with critical biological functions required for mouse development and survival. On the contrary, homozygous RanBP9-TT animals do not display any obvious phenotype and both male and female mice are fertile.

**Figure 2.**
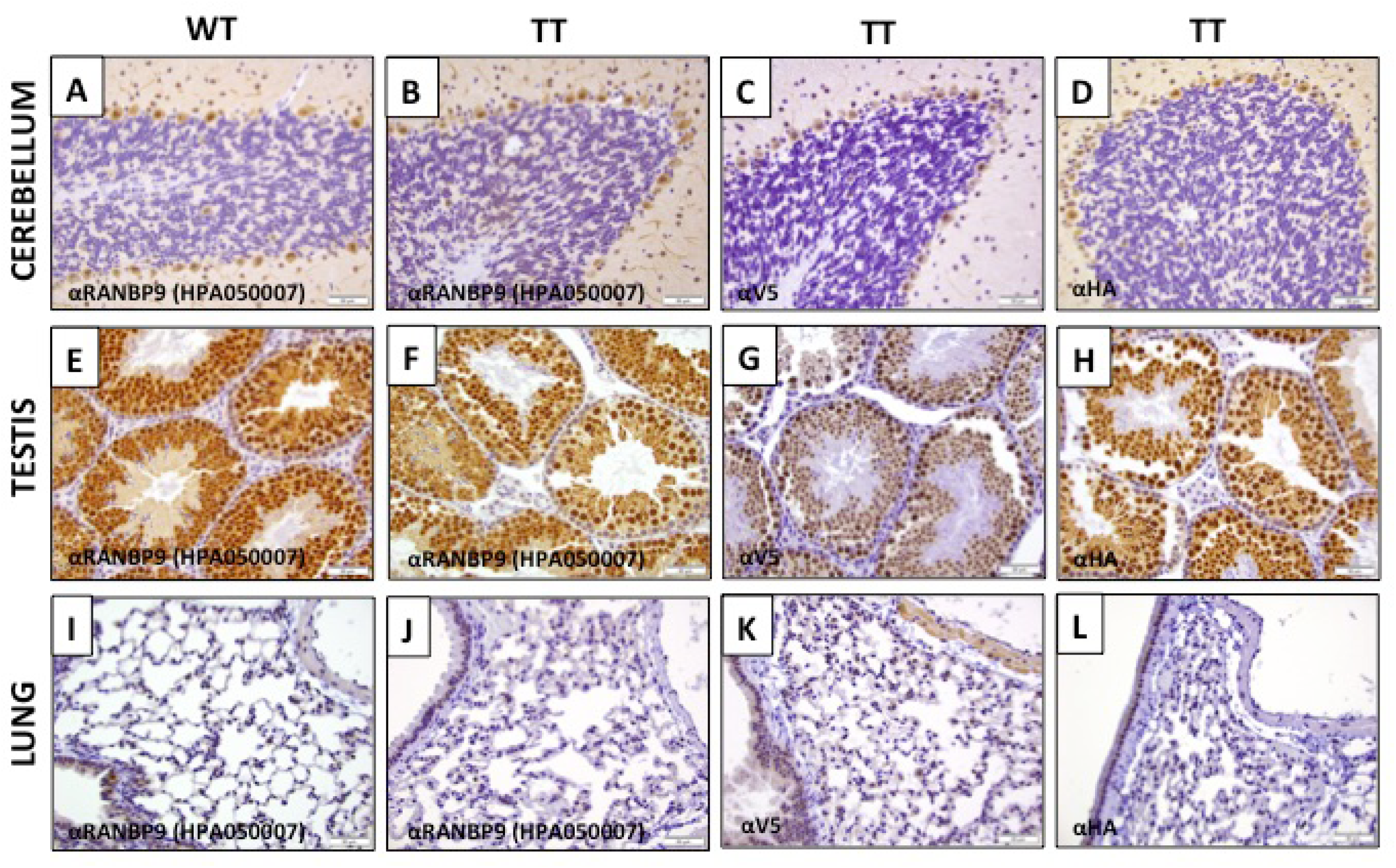
IHC detection of RanBP9 by αV5 in RanBP9-TT mice compared with detection by αRanBP9 specific antibody in RanBP9-TT and WT mice. **A**-**D**) Cerebellum; **E**-**H**) Testis; **I**-**L**) Lung. Sections from indicated organs from WT mice (**A**, **E**, **I**) and RanBP9-TT mice (**B**, **C**, **D**, **F**, **G**, **H**, **J**, **K**, **L**) were stained with αRanBP9 HPA050007 antibody (**A**, **B**, **E**, **F**, **I**, **J**), or αV5 specific antibody (**C**, **G**, **K**) or αHA specific antibody (**D**, **H**, **L**).

**Table 1.** RanBP9-TT mice are viable and fertile. Pairs of parents of the indicated genotypes were set up for breeding. In all cases the expected Mendelian ratio of homozygous and heterozygous mutant mice was obtained. The number of male and female mice obtained from the different crosses was similar. Finally, both males and females homozygous RanBP9-TT mice were able to reproduce in numbers similar to WT C57Bl/6Tac present in the colony. A statistical analysis (Chi-squared test; Graphpad Prism®) did not result in any significant difference between the expected and the observed number of animals of a specific genotype or gender.

| Female breeder genotype |  | Male breeder genotype |  | PROGENY GENOTYPE |  |  |  |
| --- | --- | --- | --- | --- | --- | --- | --- |
| HET | x | HET |  | WT | HET | HOMO KI | Tot |
|  |  |  | Females | 2 | 5 | 5 | 12 |
|  |  |  | Males | 7 | 5 | 1 | 13 |
|  |  |  | Tot | 9 | 10 | 6 | 25 |
| HOMO | x | HET |  | WT | HET | HOMO KI | Tot |
|  |  |  | Females | 0 | 8 | 1 | 9 |
|  |  |  | Males | 0 | 2 | 7 | 9 |
|  |  |  | Tot | 0 | 10 | 8 | 18 |
| HOMO | x | HOMO |  | WT | HET | HOMO KI | Tot |
|  |  |  | Females | 0 | 0 | 4 | 4 |
|  |  |  | Males | 0 | 0 | 6 | 6 |
|  |  |  | Tot | 0 | 0 | 10 | 10 |

### Addition of V5-HA double tag at the C-terminus allows faithful immunohistochemical detection of endogenous RanBP9

We then examined the expression of RanBP9 in mouse tissues by immunohistochemistry (IHC). We obtained tissue sections from RanBP9 WT and RanBP9-TT animals and stained them with αRanBP9 specific antibody (HPA050007), αV5, or αHA specific antibody. In **Figure 2**, a selected array of tissues with significant expression of RanBP9 is shown. In RanBP9-TT Cerebellum, Testis, and Lung αV5 and αHA staining are similar in pattern but sharper than the αRanBP9 specific antibody. Immunoreactivity in the cerebellum is limited to the nucleus and cytoplasm of Purkinje cells. Intense nuclear and cytoplasmic staining is also apparent within the spermatocytes and round spermatids in seminiferous tubules of the testis. In the lung, labeling is detected in the cytoplasm of alveolar macrophages and the cardiac muscle normally present within pulmonary veins. On the other hand, αtag antibodies do not detect RanBP9 in sections of the same tissues from WT animals (**Figures S2**). Finally, we also performed a more comprehensive IHC survey of several other organs/tissues using both αV5 (**Figures S3**) and αHA (**Figures S4**). Elsewhere in the nervous, immunoreactivity is localized to the nucleus and cytoplasm of neurons in the cerebrum, CA1 and CA3 regions of the hippocampus, thalamus and brainstem, ganglia throughout the body, as well as the cytoplasm of choroid plexus epithelial cells. Cardiac and skeletal muscle is consistently immunoreactive, whereas smooth muscle labeling is variable and limited to the myometrium of the oviduct, uterus, small intestine and variably within arterioles. Squamous epithelial cells in the epidermis, hair follicles, forestomach and vagina display cytoplasmic staining. Epithelia throughout an array of other tissues also exhibit cytoplasmic staining including in: sebaceous glands (sebocytes), mammary glands (ducts), endometrium (including stroma), ovary (follicular and granulosa cells), oviduct, prostate gland, glandular stomach (parietal and mucous cells), small intestine, cecum (intensely immunoreactive), colon and adrenal cortex. The nucleus of white adipocytes stain in contrast to the diffuse cytoplasmic staining in brown adipocytes. Germinal centers within the spleen and lymph nodes exhibit nuclear staining in proliferating B lymphocytes. Hematopoietic cells such as megakaryocytes within the splenic red pulp also demonstrate cytoplasmic staining. Overt labeling was not detected in the kidney, urinary bladder, liver or pancreas.

### V5-HA-tagged RanBP9 mRNA and protein expression is similar to wild type RanBP9

Next, we sought to measure the amount of RanBP9-TT protein and messenger RNA in different tissues by qRT-PCR and WB. We extracted protein of tissues/organs with various levels of reported expression ^12^. WB analysis probing with antibodies specific to RanBP9 (**Figure 3A**), or αV5 (**Figure 3B**), or αHA (**Figure 3C**) shows that the levels of expression of RanBP9-TT are similar to the levels of RanBP9 in wild type animals. Accordingly, we did not detect any significant difference between levels of RanBP9 mRNA in wild type and RanBP9-TT mice (**Figure 3D**).

**Figure 3.**
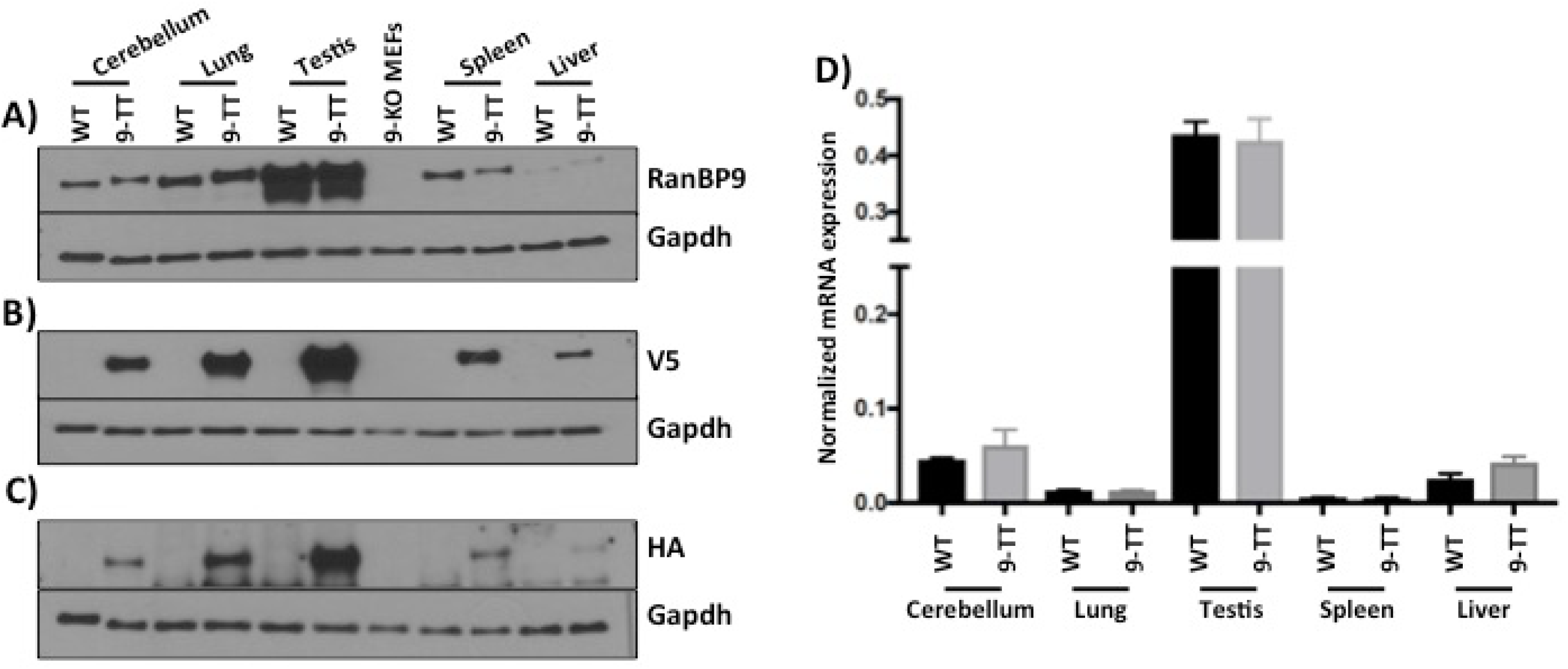
Protein and transcript expression of RanBP9 in selected adult RanBP9 wild type and RanBP9-TT mouse tissues. Proteins were extracted from mouse cerebellum, lung, testis, spleen, and liver of RanBP9 wild type (WT) and RanBP9-TT (9-TT) mice. In addition, proteins were also extracted from RanBP9-KO MEFs (9-KO MEFs) and used as negative control. **A**) Detection of RanBP9 wild type and V5-HA-tagged by αRanBP9 specific antibody. **B**) Detection of RanBP9 wild type and V5-HA-tagged by αV5 specific antibody. **C**) Detection of RanBP9 wild type and V5-HA-tagged by αHA specific antibody. **D**) Measurement of RanBP9 mRNA by RT-PCR. Expression levels of RanBP9 were assessed in Cerebellum, Lung, Testis, Spleen, and Liver from WT mice and BP9-TT mice. Expression levels in WT and TT tissues are represented in black and grey bars respectively. Each sample was analyzed in triplicate. Each well was normalized to average values of β-actin to obtain a ΔCt=2^-(FAM^ ^dye^ ^Ct-average^ ^Actb^ ^dye^ ^Ct)^. mRNA levels in each WT tissue were similar to their counterpart TT-tissues. A statistical analysis was performed using Graphpad Prism® to compare expression values in same organs from WT and TT mice. The Mann-Whitney (Wilcoxon rank sum) test was used and none of the values between WT and TT levels were significantly different.

These results show that the expression of RanBP9-TT is similar to wild type RanBP9 and there are no perturbations of expression caused by the insertion of the V5-HA tag.

### RanBP9-TT protein maintains the known RanBP9 WT interactions

RanBP9 is part of a poorly studied multi-subunit complex called the CTLH complex ^6^. Therefore, we sought to determine whether the addition of the double V5-HA tag at the C-terminus of RanBP9 interfered with its binding to the CTLH complex. To this aim, we generated homozygous RanBP9-TT mouse embryonic fibroblasts (MEFs) and performed immunoprecipitation (IP) using resin-conjugated αHA antibodies. We ran four different gels and we probed the IP fractions by WB. We were able to purchase specific antibodies for seven members of the CTLH complex. Gid8 and Muskelin are clearly present in IP fractions from RANBP9-TT but not from WT MEFs (**Figure 4A**). Similarly, Maea, Armc8 (**Figure 4B**), Wdr26 (**Figure 4C**) and Rmnd5A (**Figure 4D**) are present only in fractions pulled down by resin-conjugated αHA in the RANBP9-TT MEFs and not WT MEFs. The paralog of RanBP9, RanBP10 also is clearly present only in RANBP9-TT MEFs and not WT MEFs immunoprecipitated fractions with resin-conjugated αHA appearing as a sharp band of the expected molecular weight (**Figure 4E**). However, due to the poor sensitivity and specificity of commercially available αRanBP10 antibodies, to prove that it is RanBP10 that is immunoprecipitated and not a non-specific protein of similar size, we also extracted and immunoprecipitated lysates from RanBP10 knock-out/RanBP9-TT double mutant MEFs. Results clearly show the disappearance of the protein immunoprecipitated from RanBP10WT/RanBP9-TT MEF lysate indirectly demonstrating its specificity(**Figure 4E**).

**Figure 4.**
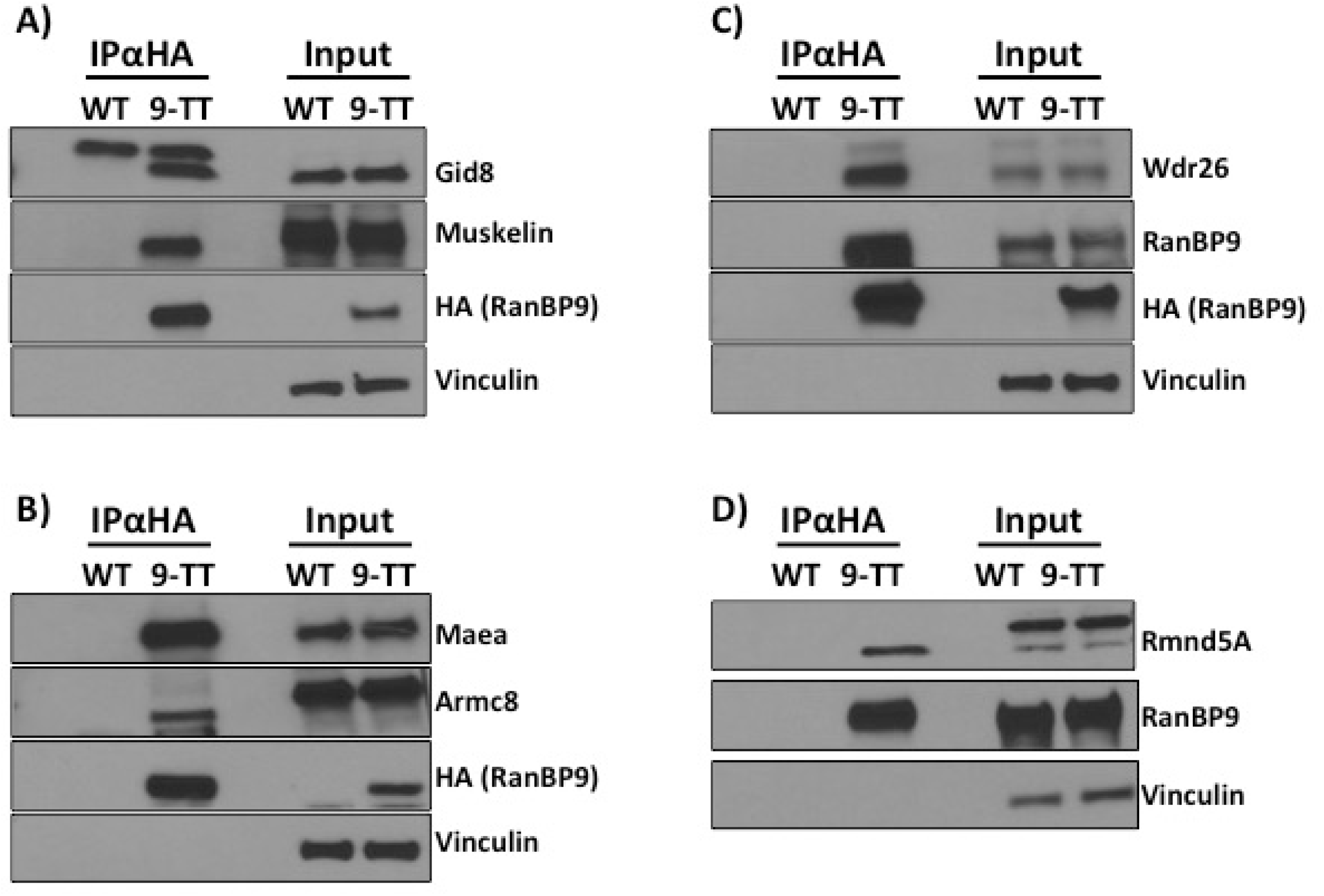
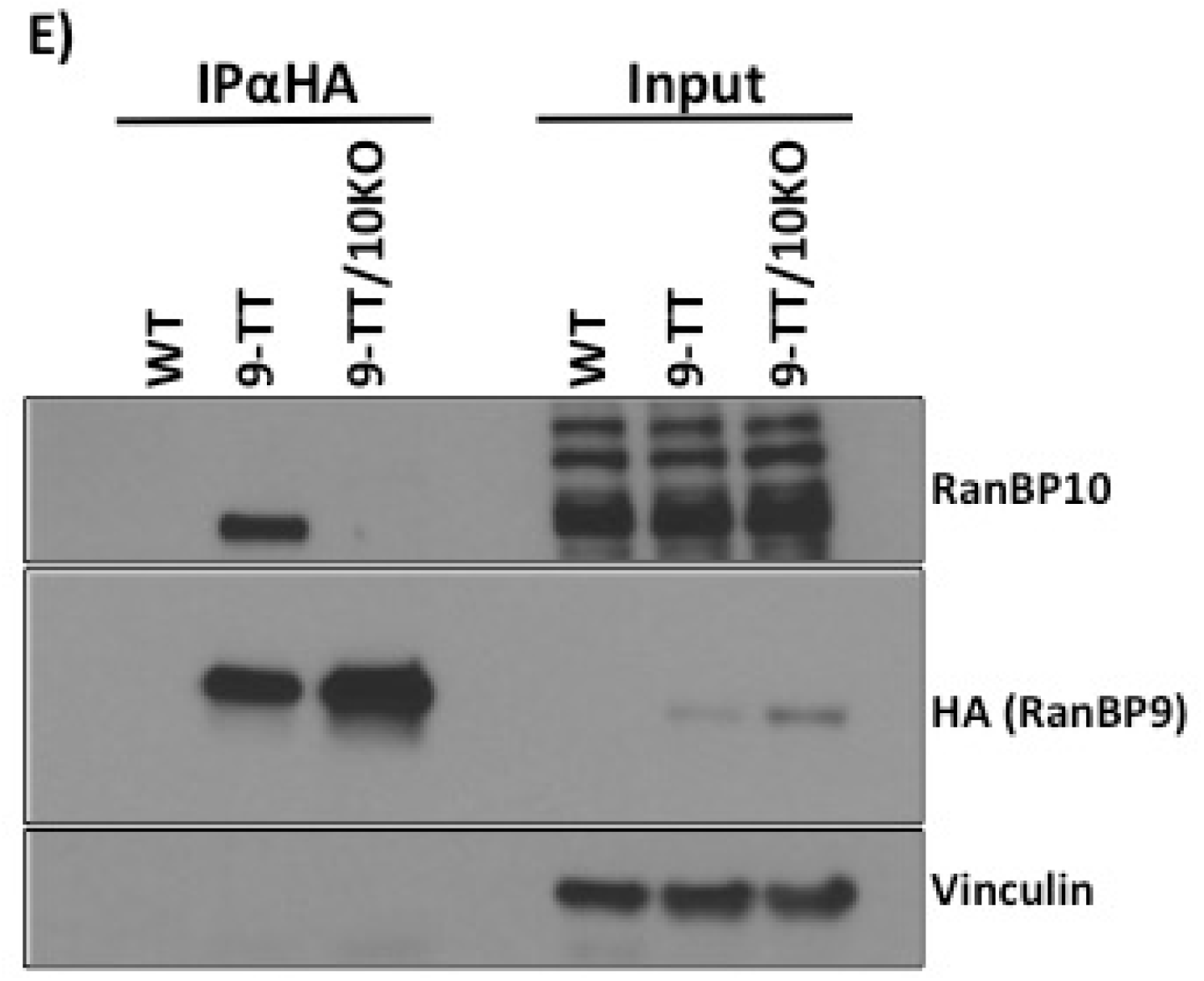
V5-HA-tagged RanBP9 maintains its ability to interact with known members of the CTLH complex. RanBP9 WT and TT Mouse Embryonic Fibroblasts (MEFs) were cultured in standard conditions and protein lysates were obtained. Resin conjugated with αHA antibodies was used to immunoprecipitate RANBP9-TT protein. IP fractions and 5% of input were run on gels to generate 4 different membranes that were probed with the indicated antibodies by WB. Vinculin is used as loading control. Shown results are representative of two independent experiments (biological replicates).

Altogether, these results clearly indicate that V5-HA-tagged RanBP9 interacts with known partners of WT RanBP9. However, since we were not able to probe by WB for the remaining members of the CTLH complex, we decided to perform a proof-of-principle small-scale IP using αV5 antibody followed by high-resolution mass spectrometry (MS) analysis (supplementary material). Again, our experiments for IP-MS/MS analysis included lysates from RanBP9 WT MEFs as negative controls. As shown in (**Table 2**), we ranked all identified proteins by number of peptide spectral matches (PSMs) or counts present in RanBP9-TT MEFs and absent in RanBP9 WT IP fractions. The exception was Muskelin, which presented one count identified in RanBP9 WT αV5 immunoprecipitates. As expected, results show that RanBP9 is robustly enriched and ranks first for number of counts in RanBP9-TT fractions while showing none in the RanBP9 WT MEF IP. Importantly, the list of proteins detected in RanBP9-TT lysates by IP-MS/MS includes the remaining 10 members of the CTLH complex within the top 19 hits (**Table 2**).

**Table 2.**
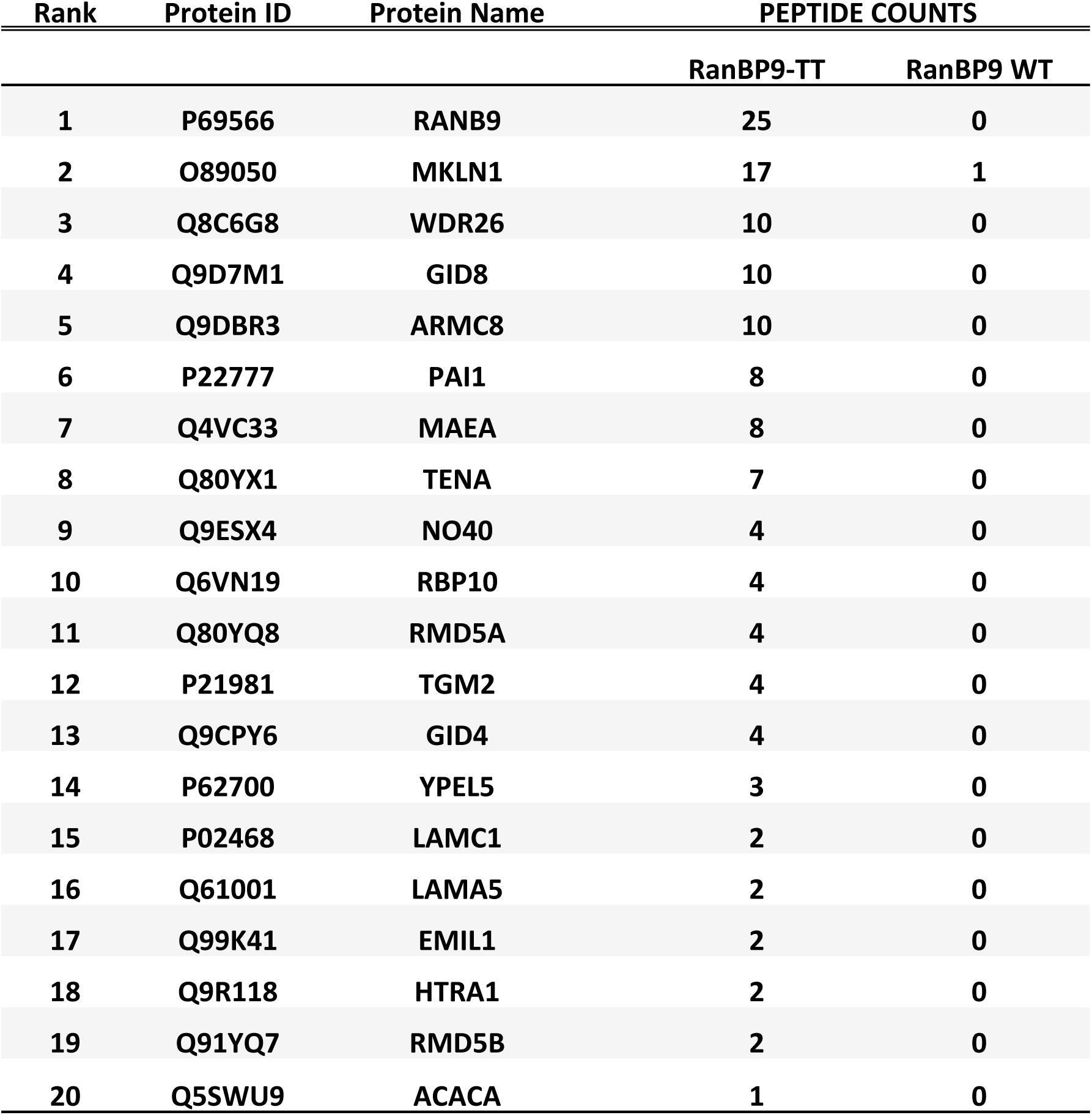
List of top 20 hits of IP-MS/MS from RanBP9-TT MEFs following ranking by number of PSMs. Protein ID, name, and number of PSMs of immunoprecipitated proteins from RANBP9-TT and WT MEFs are reported.

| Rank | Protein ID | Protein Name | PEPTIDE COUNTS |  |
| --- | --- | --- | --- | --- |
|  |  |  | RanBP9-TT | RanBP9 WT |
| 1 | P69566 | RANB9 | 25 | 0 |
| 2 | O89050 | MKLN1 | 17 | 1 |
| 3 | Q8C6G8 | WDR26 | 10 | 0 |
| 4 | Q9D7M1 | GID8 | 10 | 0 |
| 5 | Q9DBR3 | ARMC8 | 10 | 0 |
| 6 | P22777 | PAI1 | 8 | 0 |
| 7 | Q4VC33 | MAEA | 8 | 0 |
| 8 | Q80YX1 | TENA | 7 | 0 |
| 9 | Q9ESX4 | NO40 | 4 | 0 |
| 10 | Q6VN19 | RBP10 | 4 | 0 |
| 11 | Q80YQ8 | RMD5A | 4 | 0 |
| 12 | P21981 | TGM2 | 4 | 0 |
| 13 | Q9CPY6 | GID4 | 4 | 0 |
| 14 | P62700 | YPEL5 | 3 | 0 |
| 15 | P02468 | LAMC1 | 2 | 0 |
| 16 | Q61001 | LAMA5 | 2 | 0 |
| 17 | Q99K41 | EMIL1 | 2 | 0 |
| 18 | Q9R118 | HTRA1 | 2 | 0 |
| 19 | Q91YQ7 | RMD5B | 2 | 0 |
| 20 | Q5SWU9 | ACACA | 1 | 0 |

Taken all together, these results indicate that the addition of the V5-HA to RanBP9 C-terminus does not preclude its participation in the CTLH complex.

### RanBP9-TT protein interacts with Nucleolin

Previous studies have found potential interactions of RANBP9 using commercially available antibodies for IP-MS/MS approaches ^5, 6^. Most of these studies were performed in human and using different antitag antibodies. One study was performed in testes and revealed the interaction of RANBP9 with several RNA processing factors ^16^. In examining the list of proteins from our IP-MS/MS experiment, we noticed that one potential unknown interactor of RanBP9 and object of investigation in our lab could be Nucleolin (NCL). Therefore, we probed by WB lysates immunoprecipitated with αHA resin. Confirming the proteomic result, we were able to detect the presence of NCL in lysate from RanBP9-TT cells and not from RanBP9 WT MEFs (**Figure 5**). These results reveal a novel interaction between RanBP9 and NCL.

**Figure 5.**
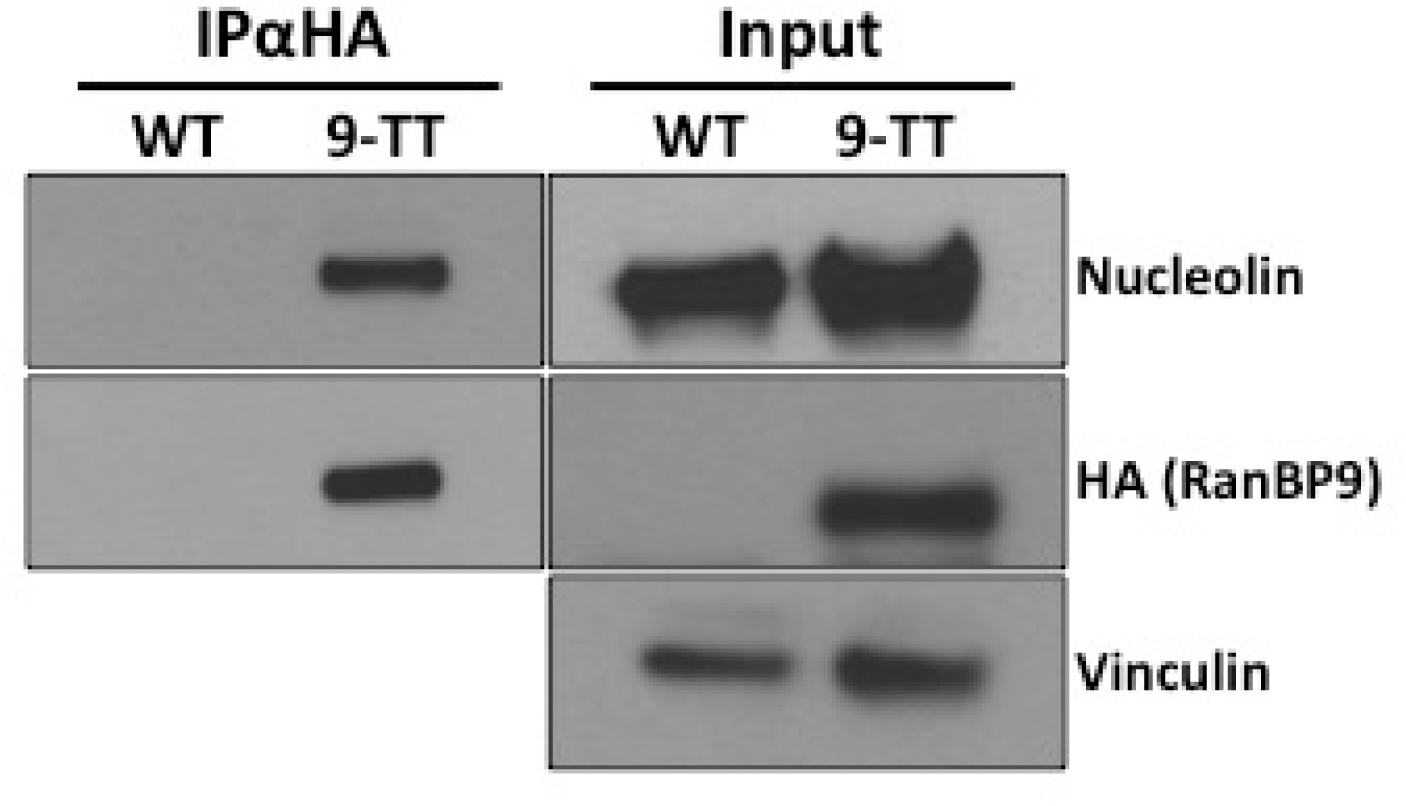
V5-HA-tagged RanBP9 interacts with Nucleolin. RanBP9 WT and TT Mouse Embryonic Fibroblasts (MEFs) were cultured in standard conditions and protein lysates were obtained. Resin conjugated with αHA antibodies was used to immunoprecipitate RANBP9-TT protein. IP fractions and 5% of input were run on gels and blotted membranes were probed with αNCL antibodies. Vinculin is used as loading control. Shown results are representative of two independent experiments (biological replicates).

## Discussion

Specific and reliable detection of proteins *in vivo* is difficult and it is estimated that researchers waste millions of dollars every year on the use of non-specific antibodies ^17, 18^. Challenges are insurmountable when the proteins of interest are not commonly studied and thus validated reagents are limited or missing. In some instances, the existence of paralogs with extensive homology further complicates the generation of tools that can unequivocally detect only the specific target. This is the case for the poorly studied scaffold protein RanBP9 and its paralog RanBP10, which share high homology in 4 out of the 5 annotated protein-protein interaction domains ^9, 10^. To date, there is only one validated antibody for detection of human RANBP9 by IHC as reported in the Human Protein Atlas project (www.proteinatlas.org). This reagent is not tested and validated for detection of mouse RanBP9 (HPA050007). In our hands, this antibody did not appear to have the sensitivity at levels similar to antibodies from other sources that performed better for biochemical detection of both human and mouse RanBP9 by conventional methods such as IP and WB. While it maybe necessary to use multiple antibodies for the detection of a target protein, this necessity generates frustration due to the quality of results and cost of reagents.

To overcome these limitations and have in hand a reliable *in vivo* tool for the study of RanBP9 in physiological and pathological paradigms, we decided to use CRISPR/Cas9 and generate RanBP9-TT mice by the addition of a double V5-HA tag at the C-terminus of RanBP9. Increasingly, biochemical tags are an excellent solution for detection problems and are widely employed for research purposes ^19^.

In order to minimize steric hindrance, we chose to use relatively small tags of 9 amino acids. In addition, we also chose to use two of them (V5 and HA) in sequence to increase the detection power, resulting in two different options.

The engineering of RANBP9-TT mice was possible through the use of CRISPR/Cas9. In fact, compared to the classical use of ES cells ^20^, in addition to being faster and efficient in various genetic backgrounds, this new technology offers the critical advantage of not leaving any “scar” in the edited allele other than the desired engineered mutation ^21, 22^. Therefore, in RanBP9-TT mice, the C-terminus of RanBP9as well as the 3’-untranslated region remained intact, preserving not only the protein structure, but also the endogenous regulation of expression (**Figure 1 and Figure S1**). The WB analysis in **Figure 3A-C** clearly shows that RanBP9 protein levels in all analyzed organs/tissues are comparable between RanBP9 WT and TT mice. Interestingly, detection of RanBP9 by the use of αV5 (**Figure 3B**) appears to have superior sensitivity and specificity (lower background) compared not only to the αHA (**Figure 3C**), but also the αRanBP9 specific antibody (**Figure 3A**). qRT-PCR measurements confirmed that there are no differences in RanBP9 mRNA levels in different organs/tissues from RanBP9-TT compared to RanBP9 WT mice (**Figure 3D**). However, it is interesting to note that levels of mRNA do not always correlate with protein levels. While testes yield the highest amount of protein and mRNA, the lungs exhibit the second highest level of protein but display a level of messenger lower than cerebellum where RanBP9 protein is significantly reduced. Similarly, liver contains the lowest amount of RanBP9 protein, but a level of mRNA higher than both spleen and lungs. These results might be explained as technical artifacts or, alternatively, suggest the existence of post-transcriptional regulatory mechanisms of gene expression.

In this regard, we have previously observed this poor correlation between levels of RanBP9 mRNA and protein in human NSCLC samples ^11^. This phenomenon leads us to believe that it is always necessary in cancer cells and tissues to measure protein levels to evaluate the expression of RanBP9 rather than relying on the amount of RNA. Therefore, a tool like the RanBP9-TT mice, which permit the unequivocal detection of RanBP9 protein, becomes invaluable for studies *in vivo*. Indeed, the IHC analysis in **Figure 2, Figure S2, Figure S3, and Figure S4** provides strong evidence that the RANBP9-TT strain can be used to clearly identify cells expressing this protein in all tissues. When crossed to *in vivo* models of different malignancies, RANBP9-TT will be of value in studying the role of this protein during tumorigenesis. One priority for our group is to show the relevance of this protein is in lung tumorigenesis and its specific expression in tumor cells versus cell populations in the microenvironment. Further, we have shown that DNA damage elicits post-translational modifications of and changes in the RANBP9 subcellular localization ^23^. Given that *in vitro* studies completely miss the complexity of organismal responses and adaptive mechanisms that ensue *in vivo*, we plan to use our new mouse model to study how the protein is modified and whether its localization changes following genotoxic stress, for example.

We have deliberately decided the placement of the TT tag based on *in vitro* evidence from our lab and other groups indicating that C-terminus tagging of RanBP9 was not impairing RANBP9 movement inside cells upon specific stimulation ^23–25^. However, such a strategy still presents the intrinsic and real risk that the added tag might block RanBP9 interactions with other proteins and interfere with its biological functions. Here, we show that vital functions and known interactions of RanBP9 are preserved in RanBP9-TT mice.

In fact, RanBP9 constitutive KO mice show a severe perinatal lethality and the occasional survivors, both males and females, are runts and sterile with gonads completely devoid of germ cells ^12, 15^. On the contrary, RanBP9-TT homozygous animals do not show any obvious phenotype. A basic observational and anatomical examination shows that TT-knock-in animals develop and grow similarly to WT littermates. Gross anatomic parameters and reproductive performance are similar in RanBP9-TT mice compared to WT littermates (**Table 1**). Specifically, RanBP9-TT mice display normal histological architecture of both testes and ovaries, among other tissues. Sperm and oocytes are present in RanBP9-TT animals without any appreciable difference compared to WT controls (**Figure 2, Figure S2, Figure S3, Figure S4**). Since sperm and oocytes are the cells in which RanBP9 is expressed at its highest levels, these results clearly show that the TT tag does not have undesired deleterious effects, nor does it interfere with fundamental biological functions of RanBP9 necessary for germ cell development and maturation ^12, 26^.

Obviously, it is impossible for us to completely exclude the possibility that the V5-HA tag is interfering at the molecular level with yet unknown biological functions of RanBP9. However, this protein is part of an evolutionary conserved and ubiquitously expressed multi-subunit E3-ligase called the CTLH (C-Terminal to LisH) complex ^4, 6–8^. Therefore, a valid, if not the best, strategy for us to test whether the TT tag interfered with molecular interactions entertained by RanBP9 was to show whether the V5-HA-tagged RanBP9 protein still participated into the CTLH complex. Here, we show by WB analysis that the immunoprecipitation of tagged RanBP9 by αHA brings down Gid8, Muskelin (**Figure 4A**), Maea, Armc8 (**Figure 4B**), Wdr26 (**Figure 4C**), Rmnd5A (**Figure 4D**), and RanBP10 and (**Figure 4E**). Identification of 7 of the 10 known CTLH members (not including RanBP9) strongly indicates that the tagged protein is incorporated into the complex and retains its activity and functionality. As reliable antibodies for the remaining CTLH members (Gid4, RanBP10, Rmnd5B, and Ypel5) are not commercially available, to further validate our findings, we performed αV5 IPs from RanBP9-TT and WT MEFs followed by tandem mass spectrometry analysis (**Table 2**). Ranking proteins by number of peptide spectral matches (PSMs) or counts in the RanBP9-TT samples and excluding those with counts in the IP from RanBP9 WT cells, all 11 members of the CTLH complex are present within the top 20 hits (**Table 2**). This proof-of-principle experiment, performed in duplicate, demonstrated that the entire complex was immunoprecipitated by V5-HA-tagged RanBP9. IP-MS/MS data corroborated the WB results (**Figure 4A-C**) and add further evidence that RanBP9-TT conserves the same interactions that the RanBP9 WT protein has.

Interestingly, proteins included in the IP-MS/MS list, included several unknown potential interactors of RanBP9. Here, we sought to confirm by IP-WB the interaction between RanBP9 and NCL (**Figure 5**). This nucleolar protein is of particular interest for several reasons. First, due to its aberrant expression NCL represents an ideal target for cancer therapy in several aggressive types of malignancies ^27, 28^. Second, NCL is an RNA-binding protein involved in processing and stabilization of coding and non-coding RNA, which would expand the potential involvement of RanBP9 in RNA metabolism. Third, NCL is involved in the response to DNA damage and might represent another unknown link between RANBP9 and p53 in cells treated with genotoxic agents. In fact, we have previously shown that when RANBP9 us ablated in NSCLC cells the levels of p53 are robustly decreased in cells treated with CDDP or IR ^11, 23^. On the other hand, NCL together with the ribosomal protein L26 bound to the 5’ untranslated region of the p53 mRNA increases the translation and consequently the levels of p53 protein ^29^. Therefore, it is tempting to speculate that in response to damage of the DNA, NCL and RANBP9 cooperate in elevating and maintaining adequate levels of p53 stabilizing its transcript. However, the mechanisms of NCL-RANBP9 interaction will need to be confirmed and studied in human NSCLC cells in the context of the DNA Damage Response. Therefore, those experiments are beyond the scope of the present investigation.

In summary, we show here that we have engineered a new mouse model bearing a tagged RanBP9 protein that retains all the known features and abilities of the endogenous wild type molecule. The TT tag allows reliable immunohistochemical detection and enrichment of RanBP9 and its known main binding partners of the CTLH macromolecular complex. This new tool will be invaluable for the study of RanBP9 functions at the organismal level a multitude of in physiological and pathological conditions and possibly reveal unknown interactions and molecular functions.

## Material and Methods

### CRISPR/Cas9 RanBP9-TT mouse generation and analysis

Mouse experiments were performed according to The Ohio State University Institutional Animal Care and Use Committee (IACUC) guidelines after a review of an institutional review board. Work performed for the present study is described in IACUC approved protocol number 2008A0009-R3 titled “Generation, analysis and training in the use of gene knock-out and transgenic rodents”; Principal Investigator: V. Coppola.

The Genetically Engineered Mouse Modeling Core of the Ohio State University Comprehensive Cancer Center generated RanBP9-TT mice by using CRISPR/Cas9 technology. Briefly, CRISPR targeting strategy was designed using the online algorithm at www.benchling.org. The synthetic single strand oligo donor DNA (ssODN) in which both V5-HA tags are contained was purchased from Integrated DNA Technologies (Coralville, Iowa, USA). Synthetic tracrRNA and crRNA was purchased from Sigma-Aldrich (Saint Louis, MO, USA). GeneArt Platinum Cas9 nuclease purified protein (cat. nr. B25642) was purchased from Invitrogen (ThermoFisher Scientific; Waltham, MA, USA). The mix of assembled sgRNA with Cas9 protein (RNP complex) and ssODN was microinjected into C57Bl/6Tac zygotes. C57Bl/6Tac WT animals were used for founder mating purposes.

### Immunohistochemical (IHC) analysis of RanBP9-TT and RanBP9 WT expression in mouse adult tissues

Adult mouse tissues were sectioned and stained by the Comparative Pathology & Mouse Phenotyping Shared Resource of the Ohio State University Comprehensive Cancer Center as previously described ^30^. Briefly, tissues were fixed in 10% (v/v) formalin, routinely processed, and embedded in paraffin for immunohistochemical characterization. Formalin-fixed, paraffin-embedded tissues were cut into 4-µm-thick sections, were dewaxed in xylene and rehydrated through graded ethanol solutions to water. Antigens were retrieved by heating the tissue sections at 96-98°C for 25 minutes in citrate solution (10 mmol/L, pH 6.0). Sections were cooled and immersed in methanol in the presence of 0.3% hydrogen peroxide for 15 minutes to block the endogenous peroxidase activity. After being rinsed first in tap water and then in PBS for 5 minutes, sections were subsequently incubated with the indicated antibodies (rabbit αRanBP9 antibody, 1:50, Sigma-Aldrich, cat. nr. HPA050007; goat αV5 antibody, 1:350, Abcam cat. nr. ab95038; and, rabbit αHA, 1:800, Cell Signaling cat. nr. C29F4) at 4°C overnight. As negative controls, slides were incubated with rabbit or goat IgG instead of the primary antibody. The sections were rinsed with buffer and then incubated with horseradish peroxidase-labeled secondary antibody (RanBP9: αrabbit, 1:1000; V5: αgoat, 1:500; HA: αrabbit, 1:500) for 30 minutes. 3,3’-Diaminobenzidine (DAB) was used as the chromogen, and hematoxylin as the nuclear counterstain. Sections were then dehydrated, cleared and mounted permanently with glass coverslips. Slides were photographed with an Olympus BX45 light microscope with attached DP27 digital camera and corresponding CellSen software (B&B Microscopes Limited, Pittsburgh, PA).

### WB analysis of RANBP9 expression

Mouse tissues were homogenized on ice in NP-40 buffer supplemented with Halt Protease and Phosphatase Inhibitor cocktail (Thermo Fisher Scientific cat. nr. 78442). Protein concentration was determined using the Bio-Rad protein assay dye (Bio-Rad cat. nr. 5000006). Western blot analysis was performed using 30 to 50 µg proteins run on Mini-PROTEAN TGX precast gels (Bio-Rad cat. nr. 456-1094). αRANBP9 primary antibody was purchased from Abcam (cat. nr. ab140627) and used at a dilution of 1:2,000 in 5% non-fat milk in TBS-T. Signals were detected with HRP-conjugated secondary antibodies (GE Healthcare) and the chemiluminescence substrate Supersignal wet pico PLUS (Thermo Fisher Scientific). Equivalent loading among samples was confirmed with αGAPDH (Cell Signaling cat. nr. 3683S).

### RT-PCR analysis of RanBP9 expression

Murine tissues were homogenized in ice using 1 ml TRIzol Reagent (Life technologies cat. nr. 15596018) following the manufacturer’s protocol. The extracted RNA was quantified using Nano-Drop 2000. To analyze RanBP9 gene expression, real time quantitative polymerase chain reaction (RT-qPCR) was performed from complementary DNA (cDNA). Total RNA (500 ng) was reverse transcribed using the High Capacity cDNA Reverse Transcription Kit (Applied Biosystem cat. nr. 4368814) following the manufacturer’s instructions. The cDNA was used for quantitative real time PCR with TaqMan fast advanced master mix (Applied Biosystems cat. nr. 4444557). Samples were amplified simultaneously in triplicates in one assay run. Analysis was performed by Prism 7.0 software (Graphpad Prism®) using the Δ-ct method (Applied Biosystems). The RANBP9 probe used was Hs00170646_m1.

### Immunoprecipitation

RanBP9 was immunoprecipitated from total protein extracts using monoclonal αHA-Agarose antibody (Sigma cat. nr. A2095-1ML). Briefly, 1mg of total protein extracts were pre-cleared by incubating with Pierce™ Protein A/G Plus Agarose (Thermo Scientific cat. nr. 20423) for 1 hr at 4°C. Lysates were centrifuged at 4°C; the supernatant was collected and incubated with 5 μg of the primary antibody overnight at 4°C. Proteins interacting with RanBP9 were detected in the immunoprecipitates of RanBP9 using wet-transfer WB. Interactors of the CTLH complex were detected using the following antibodies: αRMND5A (Novus Biologicals cat. nr. NBP1-92337), αWDR26 (Abcam cat. nr. ab85962), αARMC8 (Proteintech cat. nr. 12653-1-AP), αMuskelin (Proteintech cat. nr. 14735-1-AP), αC20orf11 (a.k.a. GID8; Proteintech cat. nr. 24479-1-AP), αMAEA (R&D Systems cat. nr. AF7288), αRanBP10 (Millipore SIGMA; cat. nr. SAB3500163), and αNucleolin (D4C7O; Cell Signaling; rabbit mAB cat. nr. 14574).

### Immunoprecipitation Tandem Mass Spectrometry (IP-MS/MS)

Cells were separated into cytoplasmic and nuclear fractions prior to immunoprecipitation as described in Mohammed *et al* ^31^. Briefly, αV5 antibody (5 μg, Invitrogen cat. nr. R96025) was incubated with Pierce™ Protein A/G Magnetic Beads (Thermo Scientific cat. nr. 88803) at 4°C with rotation overnight. Following washes to remove unbound antibody, 1 mg of cytoplasmic or nuclear protein extract was incubated with the bead-antibody mixture at 4°C with rotation overnight. After stringent washes to remove non-specific proteins, beads were equilibrated in 100 mM ammonium bicarbonate with 3 washes prior to on-bead digestion with trypsin (800 ng; Promega cat. nr. V5280) overnight at 37°C 800 rpm. Digestion continued the next day for 4 hours after samples were supplemented with an additional bolus of trypsin (800 ng). Beads were placed in a magnetic stand to collect supernatants and dried down in a Speedvac concentrator.

Prior to mass spectrometry analysis, peptides were resuspended in loading buffer (2% ACN, 0.003% TFA) and quantified via Nanodrop. Chromatography separation and mass spectrometry methods were performed as described in Scheltema *et al* for data dependent acquisition on a Q-Exactive HF (Thermo Fisher Scientific) ^32^. Tryptic peptides (1000 ng) were isocratically loaded onto a PepMap C18 trap column (300µm x 5mm, 100Å, 5µm) at 5µL min^-1^ and analytical reversed phase separations were performed on a Dionex UltiMate 3000 RSLCnano HPLC system coupled to an EASYSpray PepMap C18 column (15cm x 50µm ID, 100Å, 2µm) over a 90 min gradient at 300nL min^-1^. RAW mass spectrometry files were converted to mzml and searched against a complete, reviewed mouse Uniprot database containing common contaminants (downloaded 04/10/2019) via OpenMS (version 2.3.0) with X! TANDEM (release 2015.12.15.2) and MS-GF+ (release v2018.01.20).

## Acknowledgements

The authors are grateful to Ms. Lu Ming from the Genetically Engineered Mouse Modeling Core of the Ohio State University Comprehensive Cancer Center for her support in mouse colony breeding as well as Ms. Juliann Rectenwald from the Comparative Pathology and Mouse Phenotyping Shared Resource of the Ohio State University Comprehensive Cancer Center for mouse necropsy and tissue preparation. We also thank Dr. Rhonda Pitsh from the Air Force Research Laboratory for assistance with MS/MS-IP.

## Author Contribution

S. S. and A.E.S. performed experiments, elaborated data, prepared figures and tables; M.G. and S.W.H. performed proteomic experiments and data elaboration; C.C.B. performed histo experiments and prepared figures; F.A. designed CRISPR/Cas9 targeting, performed experiments and prepared figures; A.O. and M.C. performed experiments; A.T. read and edited the manuscript; J.A.M. supervised proteomic experiments; R.V. read and edited the manuscript; M.A.F. supervised proteomic experiments; K.M.D.LP. supervised histo experiments, prepared figures, read and edited the manuscript; D.P. and V.C. conceived research, wrote manuscript, prepared figures and tables. All authors approved the manuscript.

## Additional Information

A.T. is a recipient of a post-doctoral fellow Pelotonia Award. M.G is a recipient of graduate student Pelotonia Award. Research reported herein was supported by The Ohio State Comprehensive Cancer Center and the National Institutes of Health under grant number P30 CA016058.

## Supplementary Material

## Supplementary Figures

**Figure S1.**
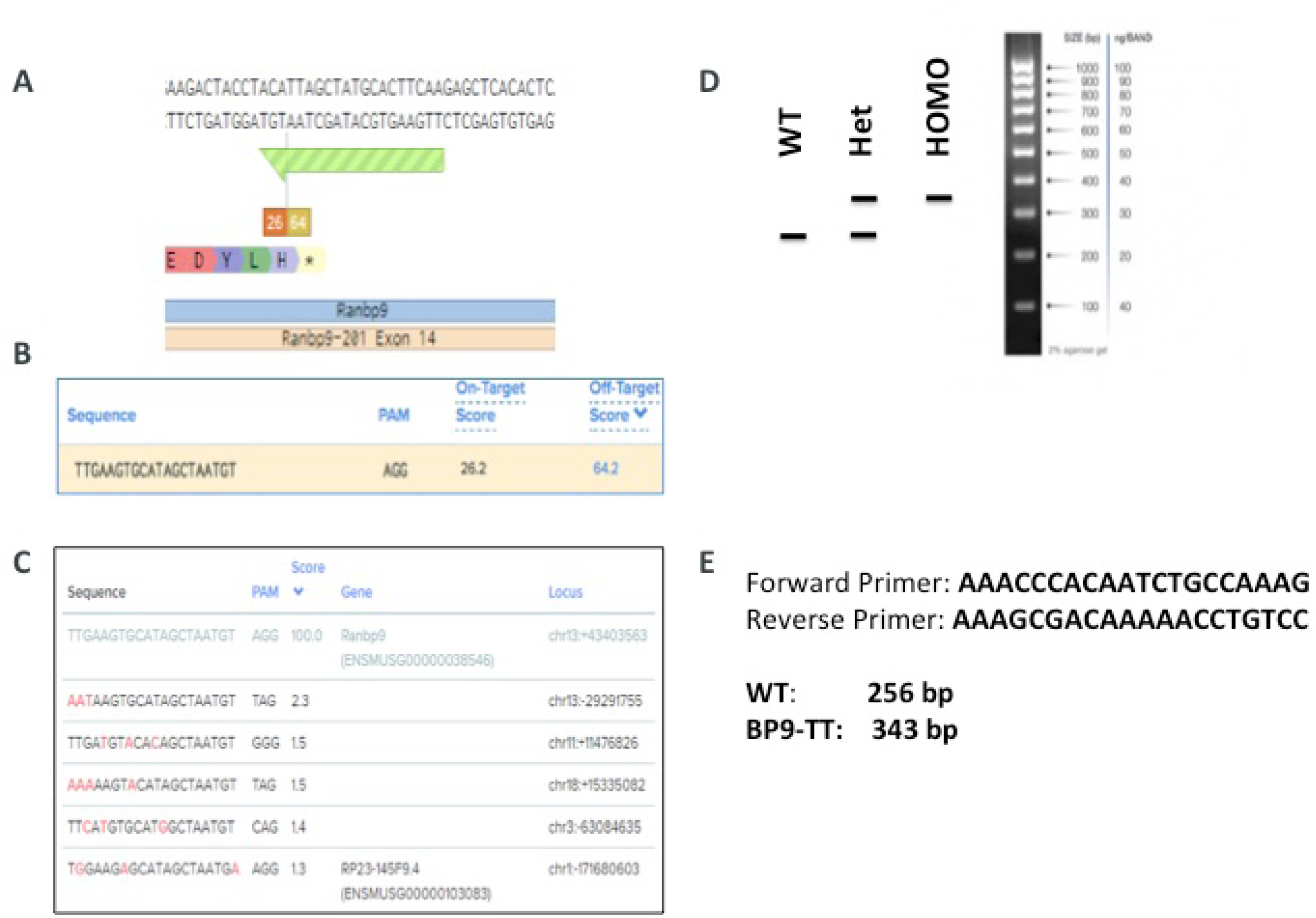
Generation of the RanBP9-TT mouse model by CRISPR/Cas9. **A**) Depiction of guide RNA (sgRNA; light GREEN) selected to target the last codon of RanBP9 and relative on-target (MAROON) and off-target (LIGHT BROWN) scores (Benchling). **B**) sgRNA sequence and scores (Benchling). **C**) Predicted off-target sites and scores using the selected sgRNA (Benchling). **D**) Virtual PCR prediction for screening of positively recombined RanBP9-TT KI animals. **E**) Sequence of primers used for mutant KI screening and expected size (base pair) of amplicons for WT and RanBP9-TT animals.

**Figure S2.**
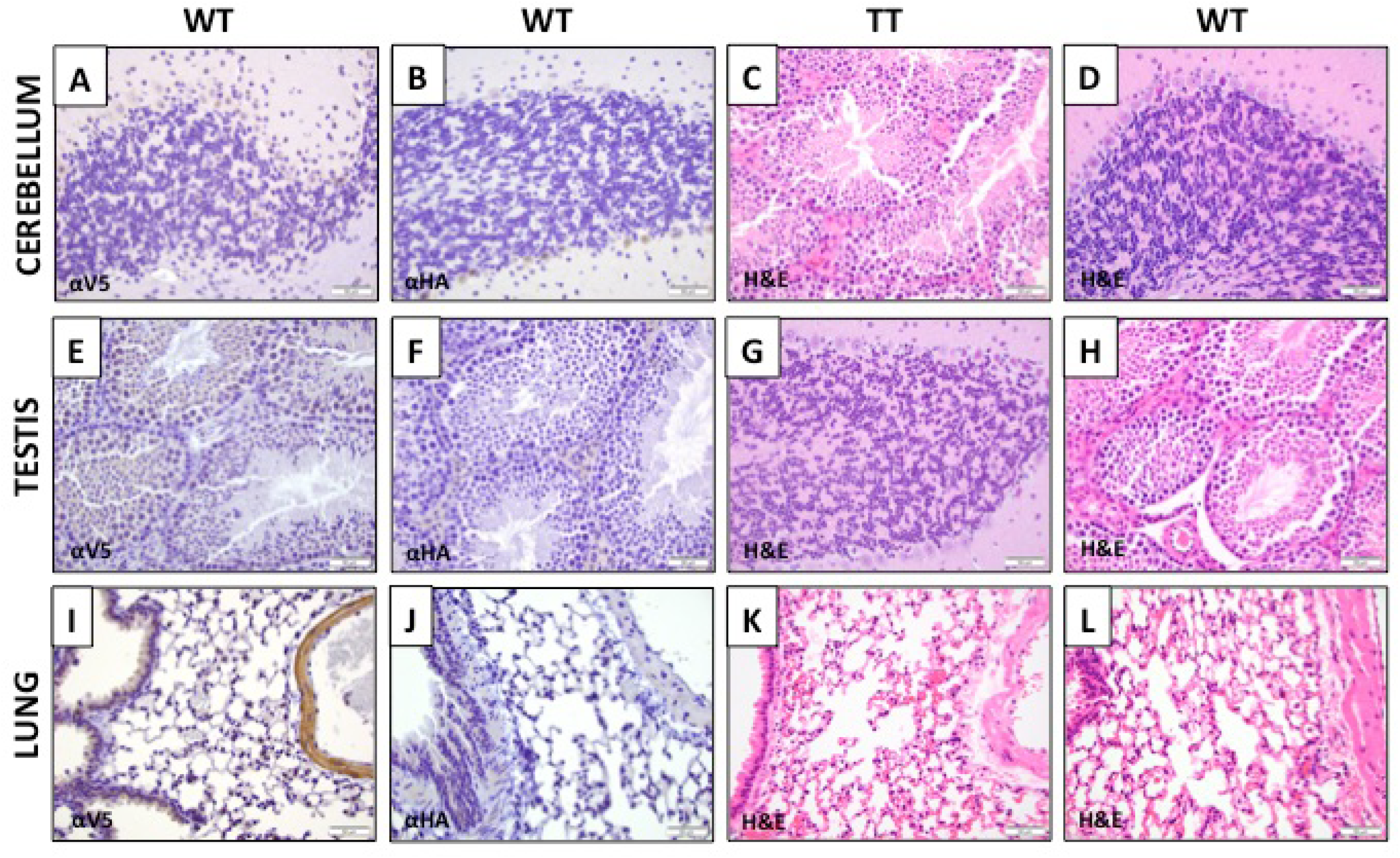
IHC detection of RanBP9 by αV5 or αHA in homozygous RanBP9-TT mice compared with detection by αRanBP9 specific antibody. Sections from same organs shown in Figure 2. **A**-**D**) Cerebellum; **E**-**H**) Testis; **I**-**L**) Lung. Sections from indicated organs from WT (**A**, **B**, **D**, **E**, **F**, **H**, **I**, **J**, **L**) and RanBP9-TT mice (**C**, **G**, **K**) were stained with αV5 (**A**, **E**, **I**), αHA (**B**, **F**, **J**) or with Hematoxylin and Eosin (H&E) (**C**, **D**, **G**, **H**, **K**, **L**).

**Figure S3.**
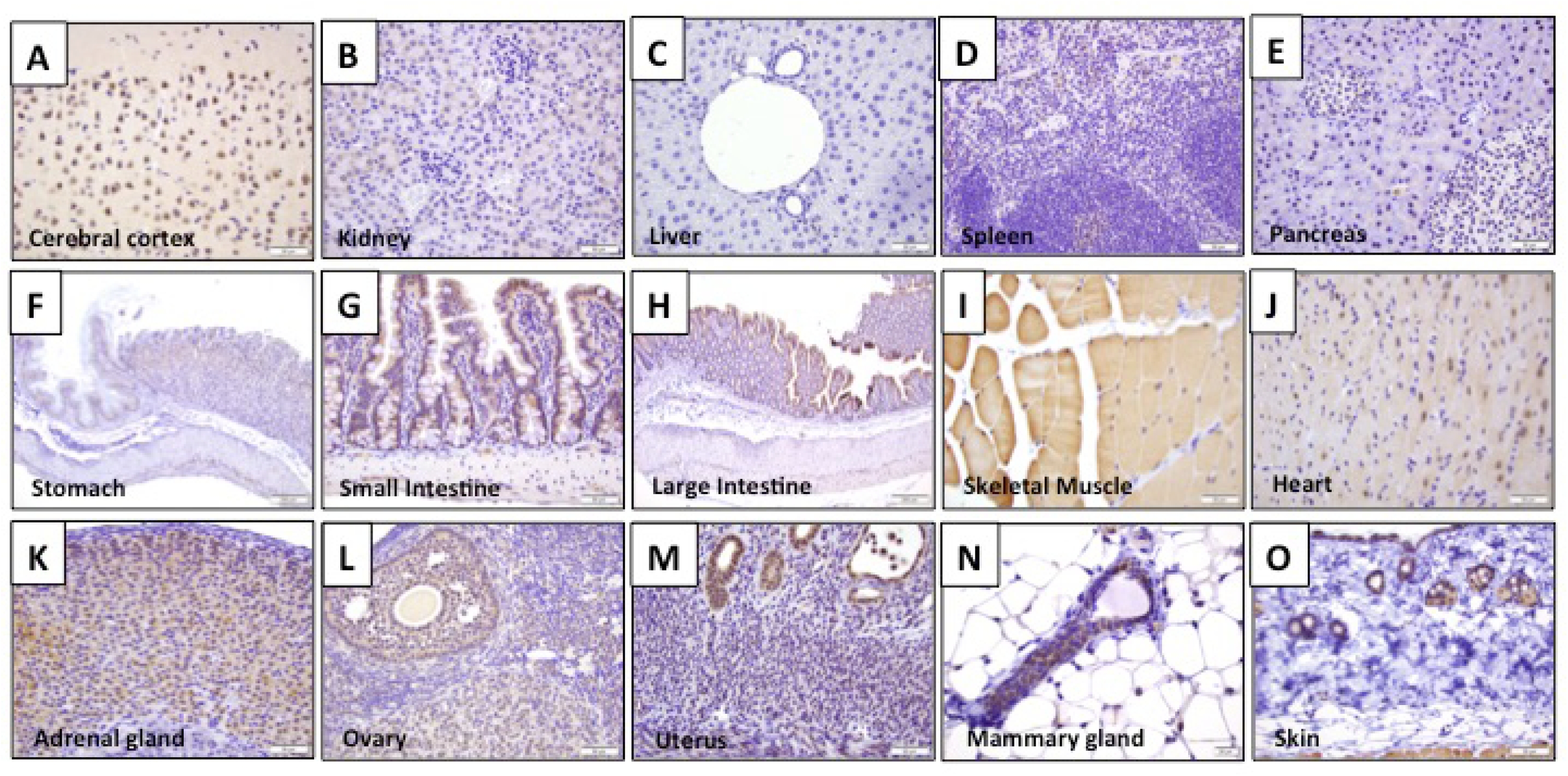
IHC detection of V5 in RanBP9-TT mice. **A**-**O**) Selection of indicated organs/tissues from RanBP9-TT homozygous mice stained with αV5 specific antibody. All pictures were taken with 40x objective and 10x eyepiece (400x). Scale bar = 50um EXCEPT: stomach (10x = 100x, scale bar = 200um); large intestine (10x = 100x, scale bar = 200um); mammary gland (60x = 600x, scale bar = 20um).

**Figure S4.**
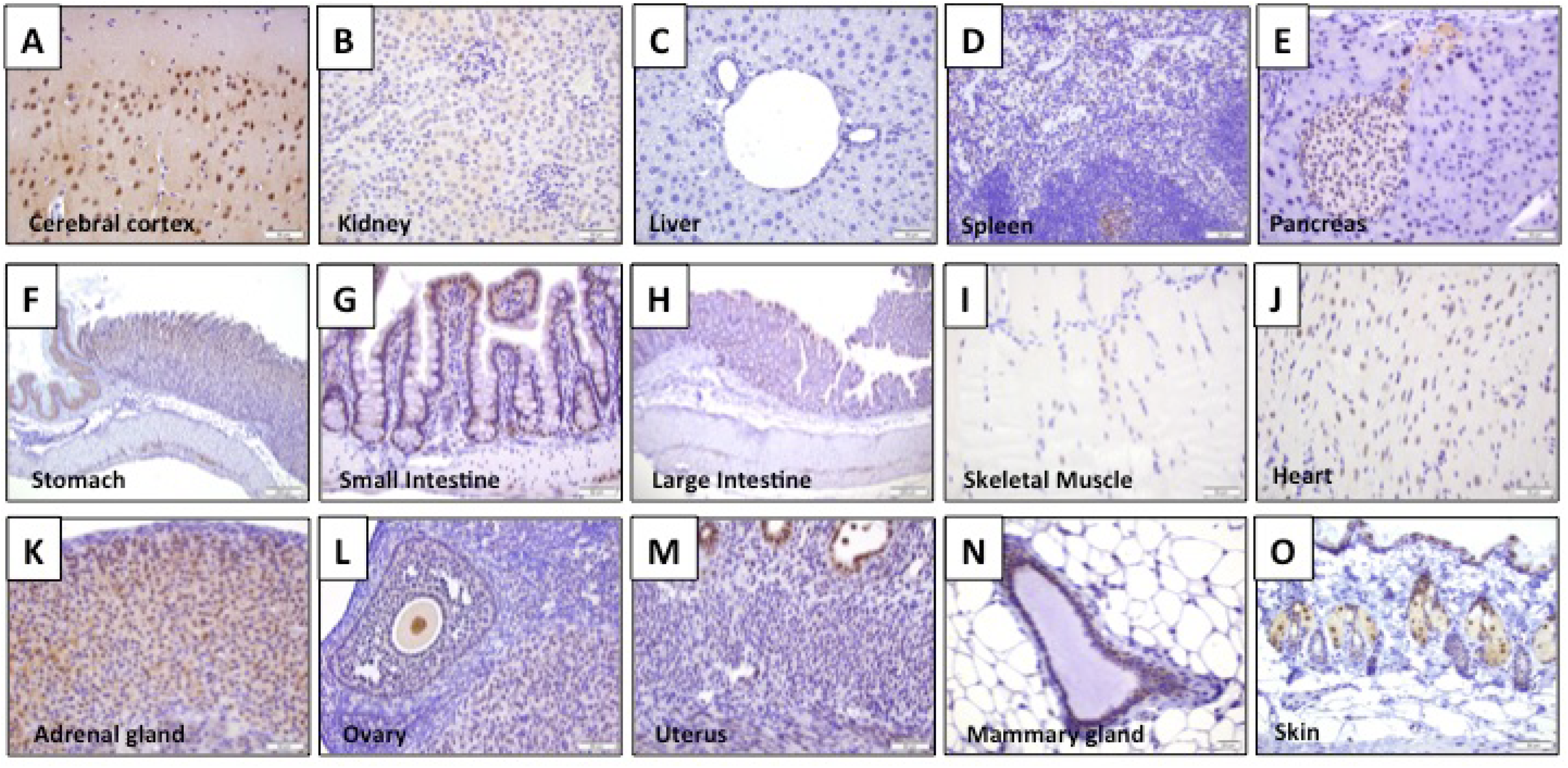
IHC detection of HA in RanBP9-TT mice. **A**-**O**) Selection of indicated organs/tissues from homozygous RanBP9-TT mice stained with αHA specific antibody. All pictures were taken with 40x objective and 10x eyepiece (400x). Scale bar = 50um EXCEPT: stomach (10x = 100x, scale bar = 200um); large intestine (10x = 100x, scale bar = 200um); mammary gland (60x = 600x, scale bar = 20um).

**Proteomic data**. The datasets generated and analyzed for this study are available in the MassIVE repository [MSV000084462] at: (ftp://massive.ucsd.edu/MSV000084462/).

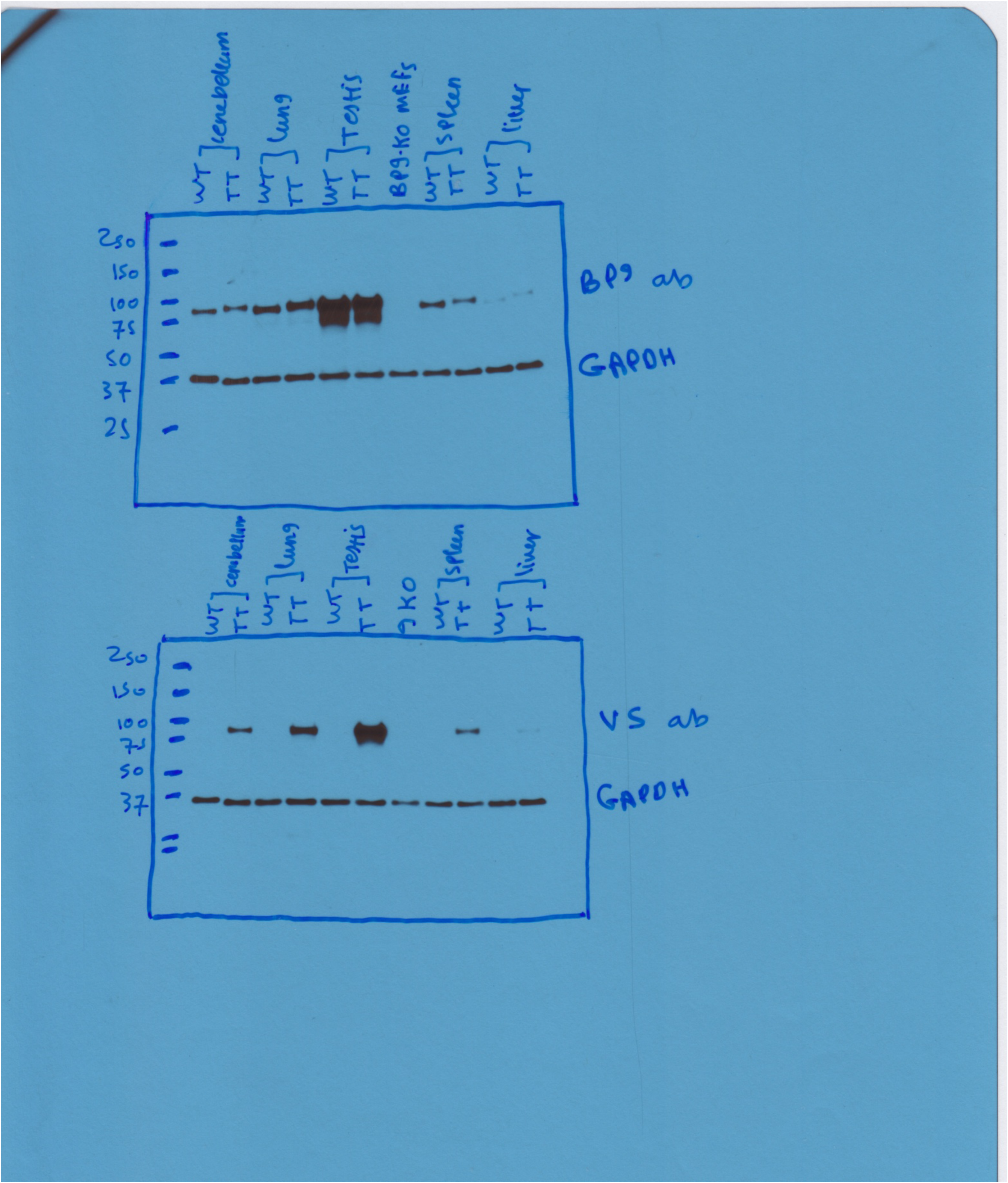
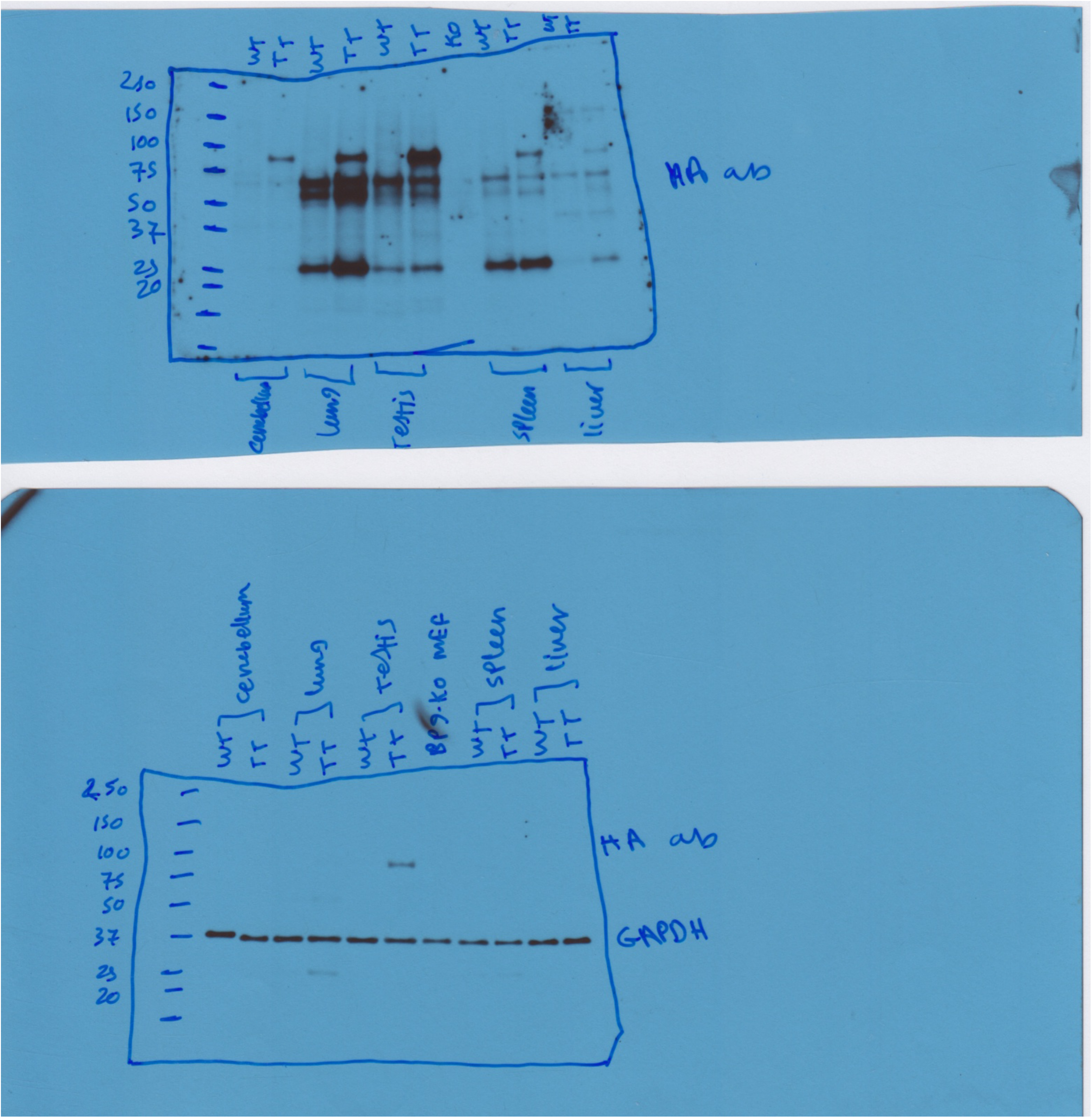
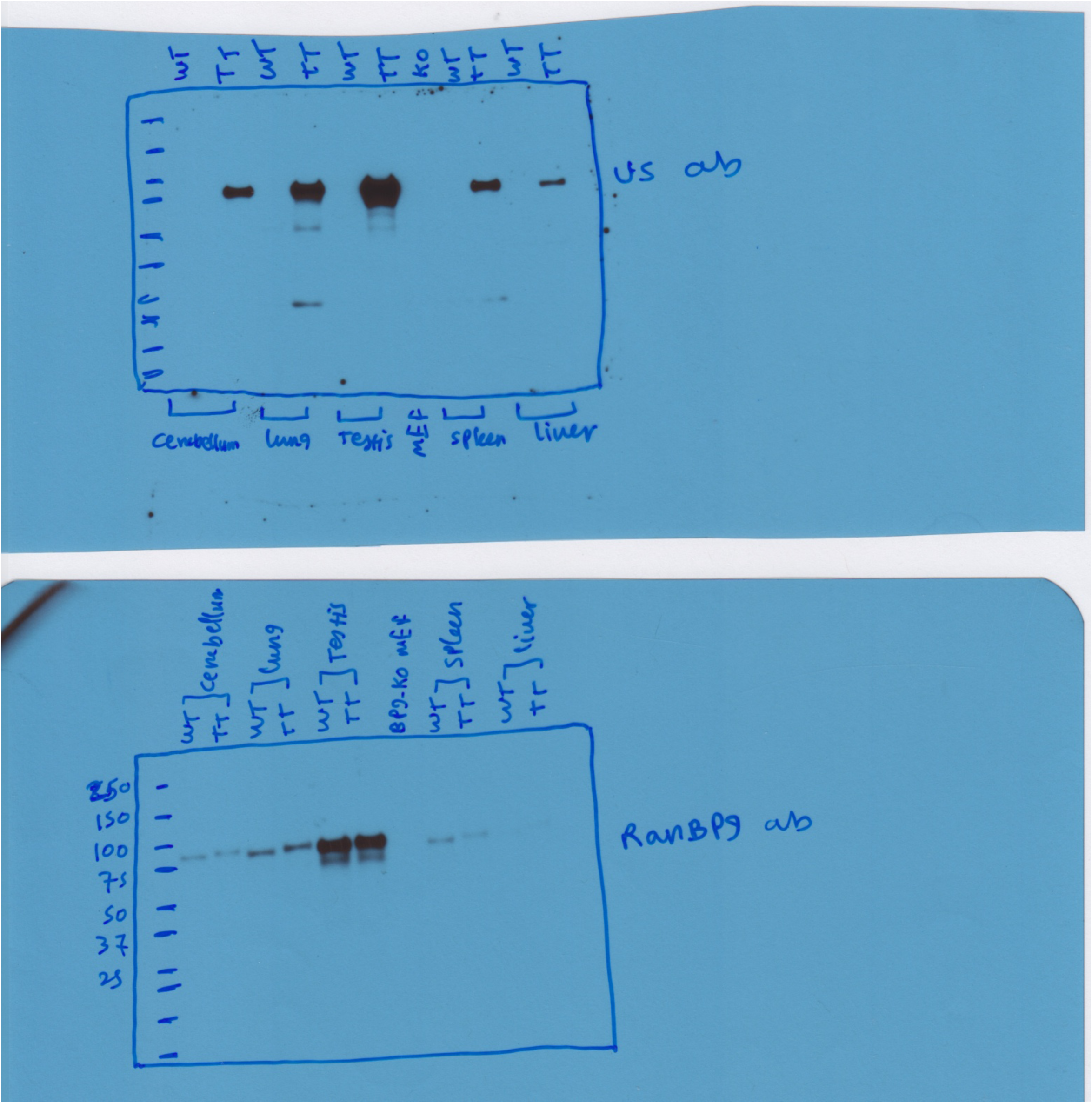
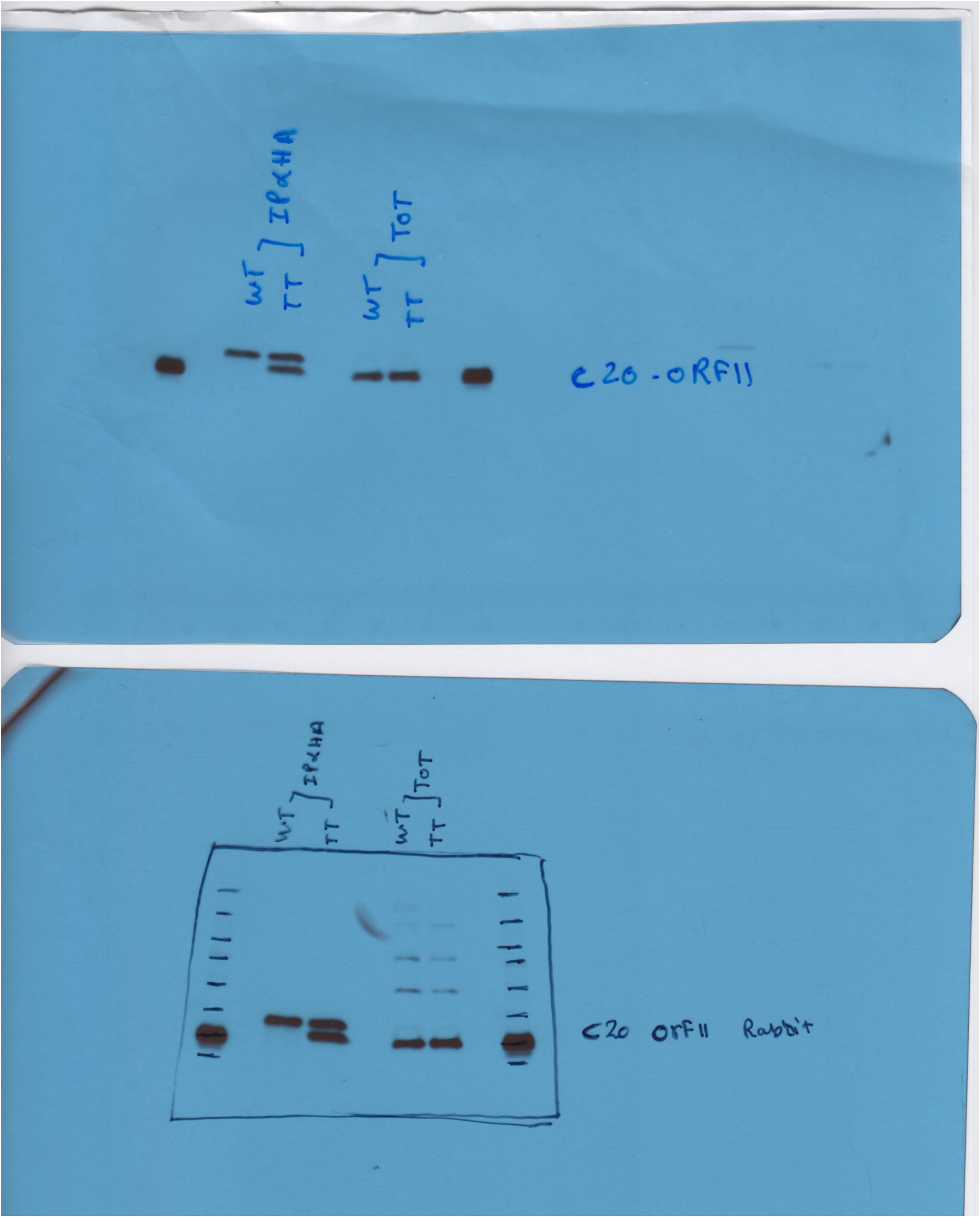
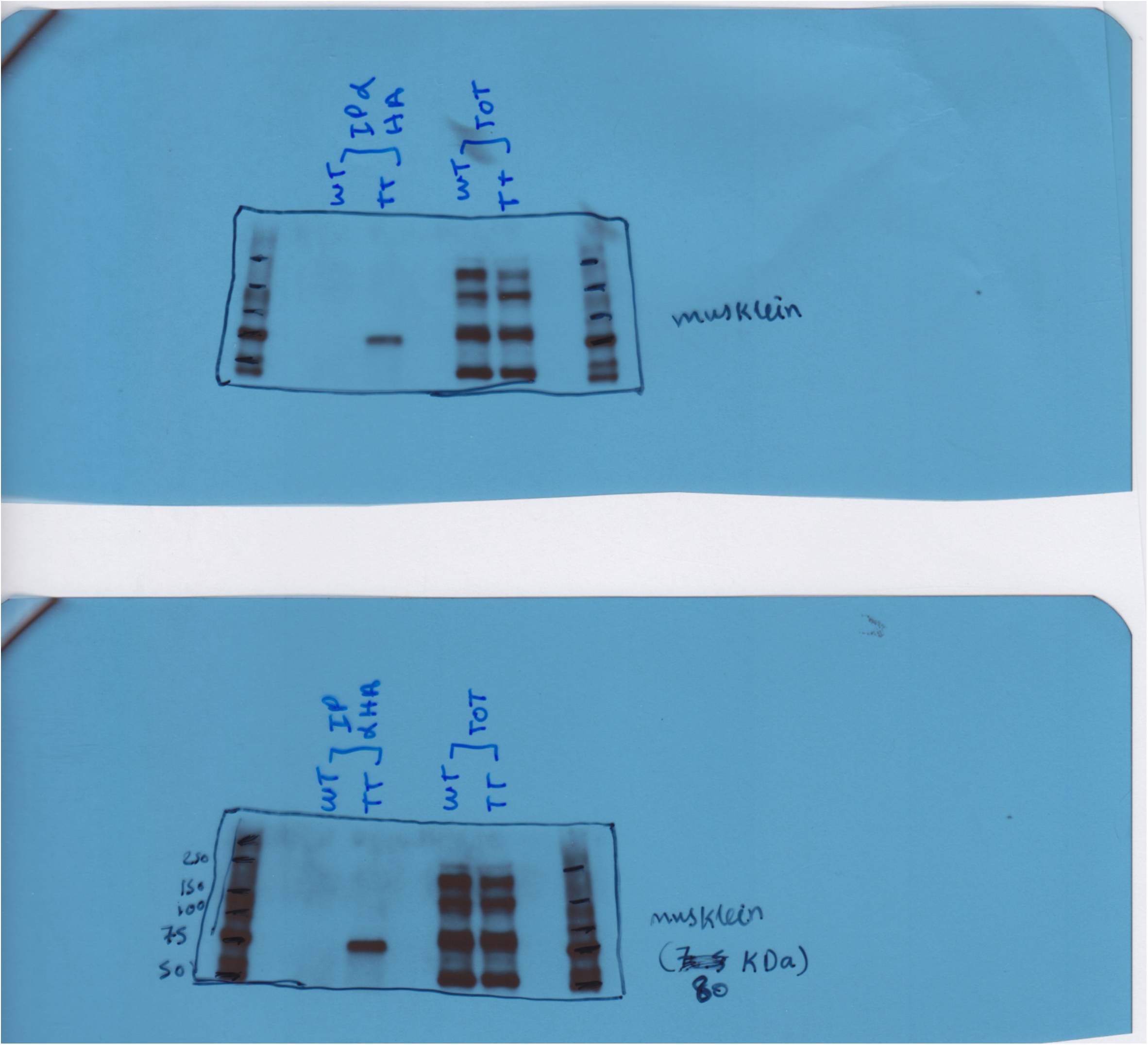
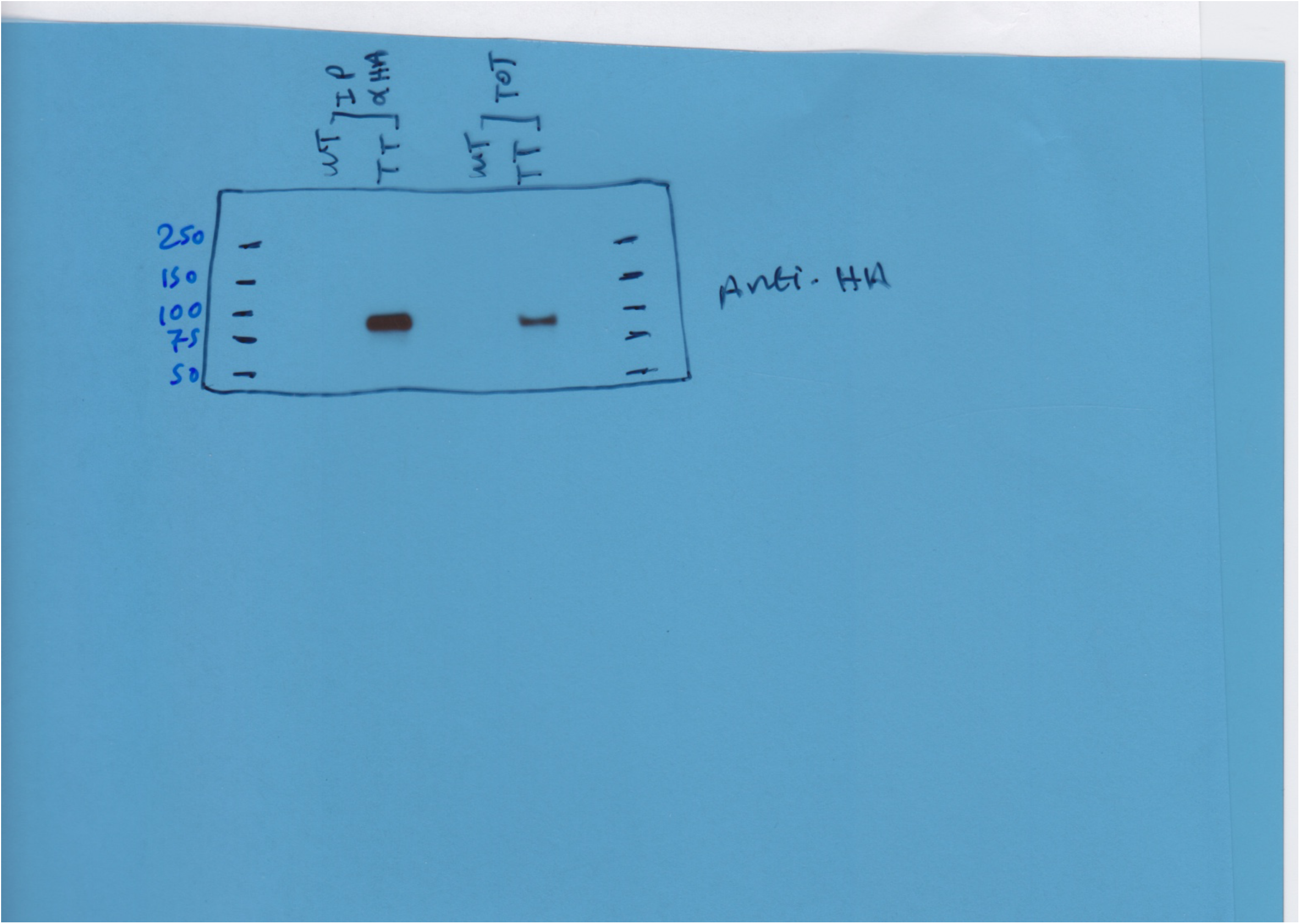
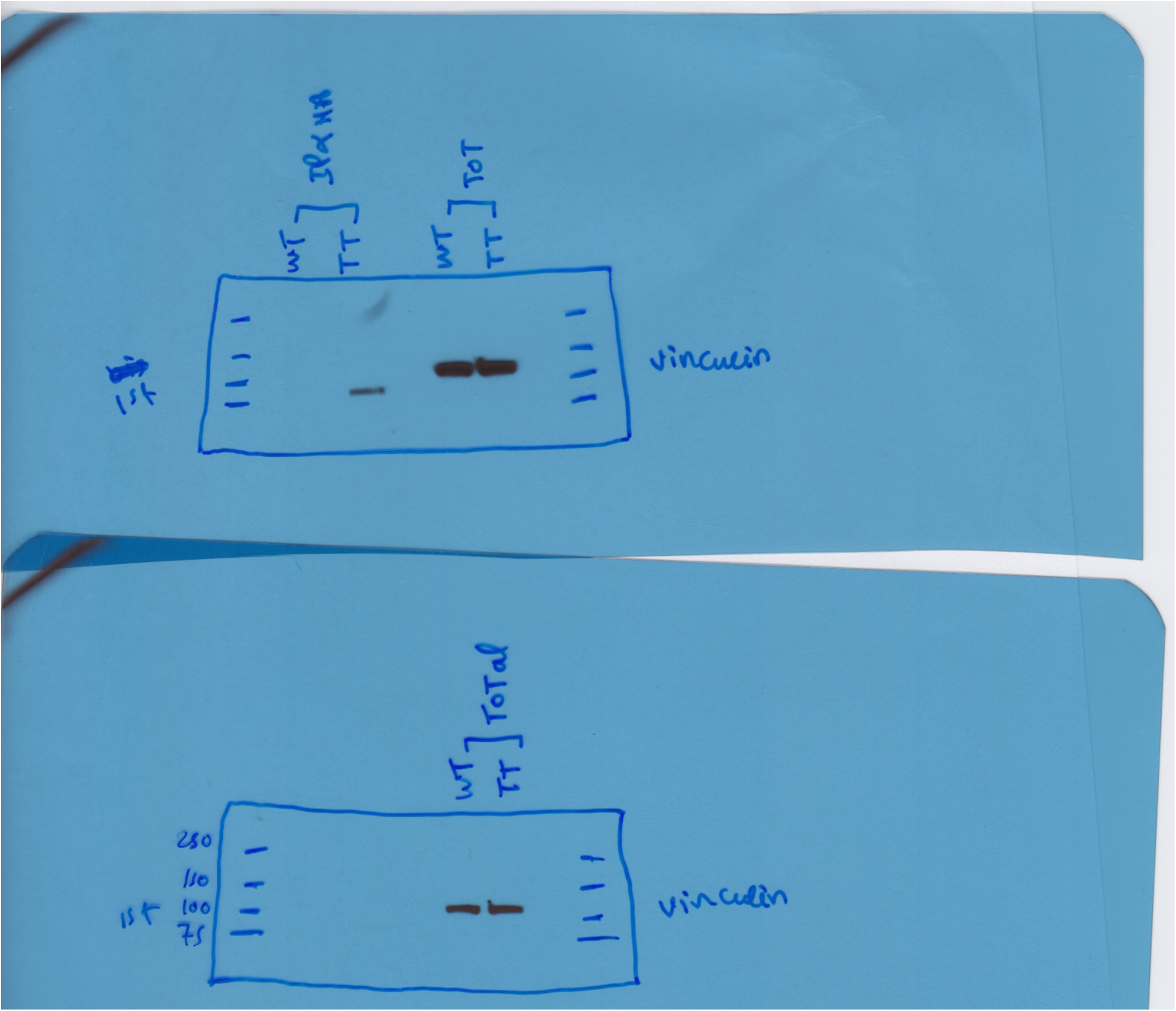
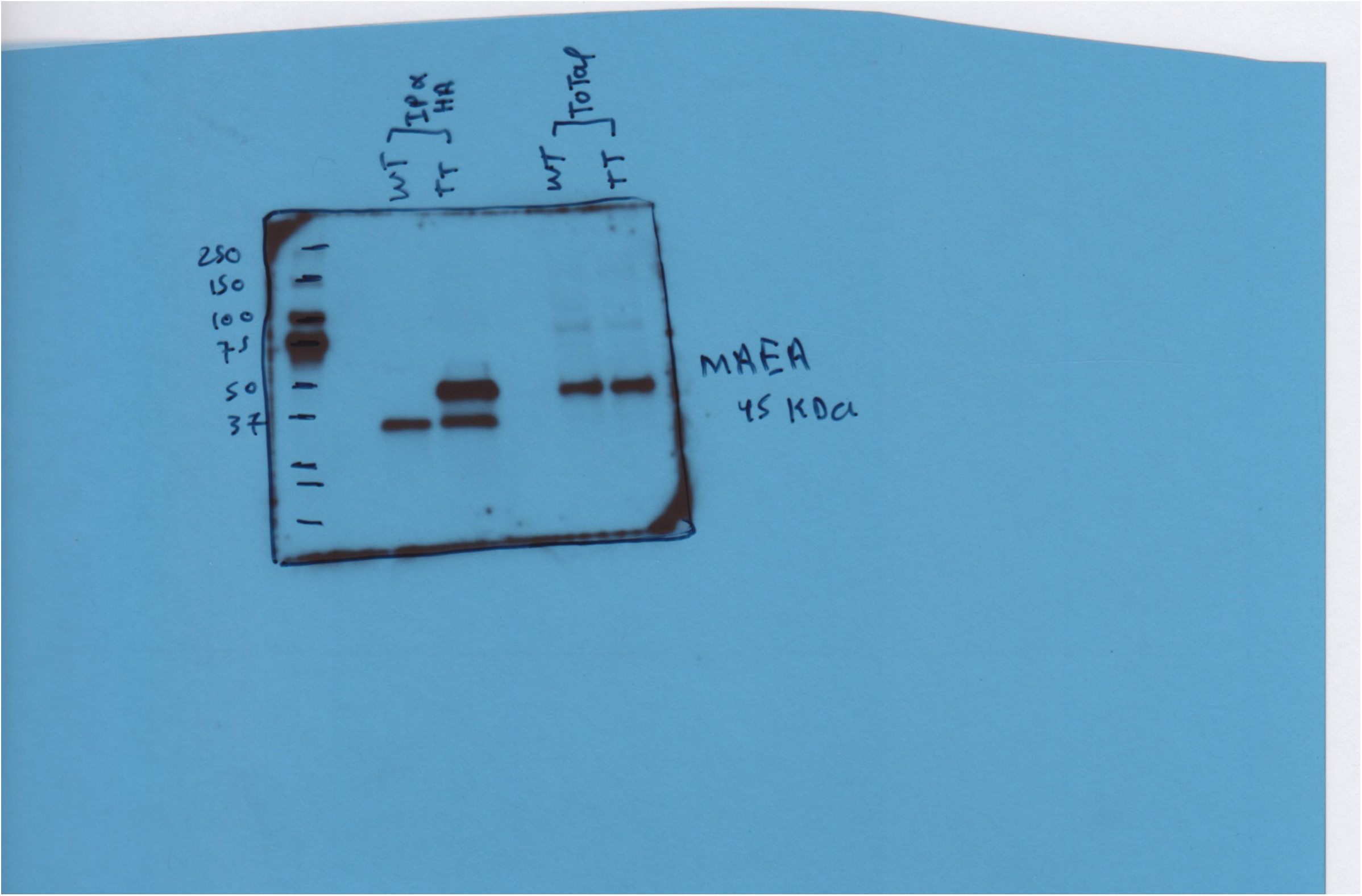
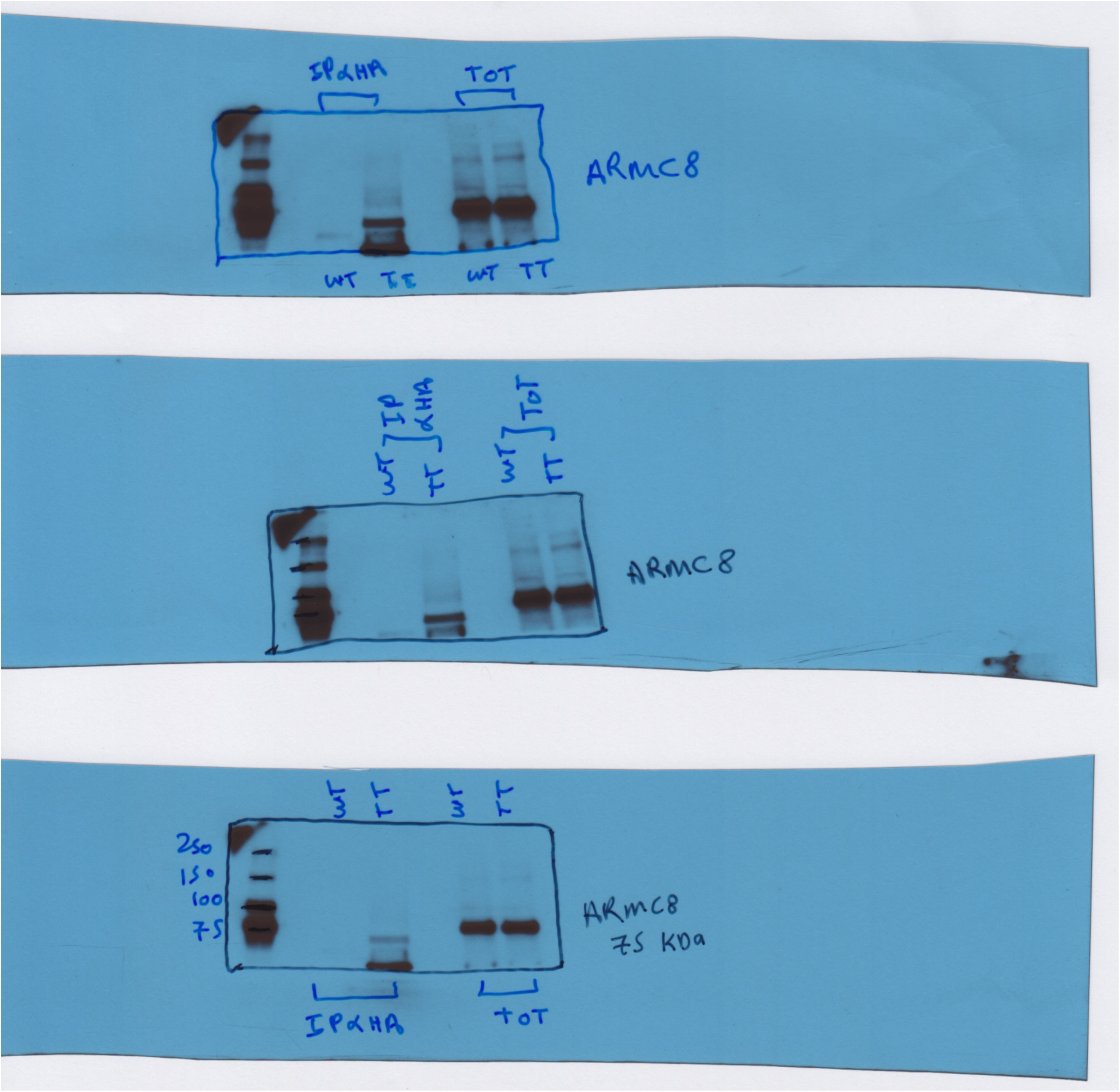
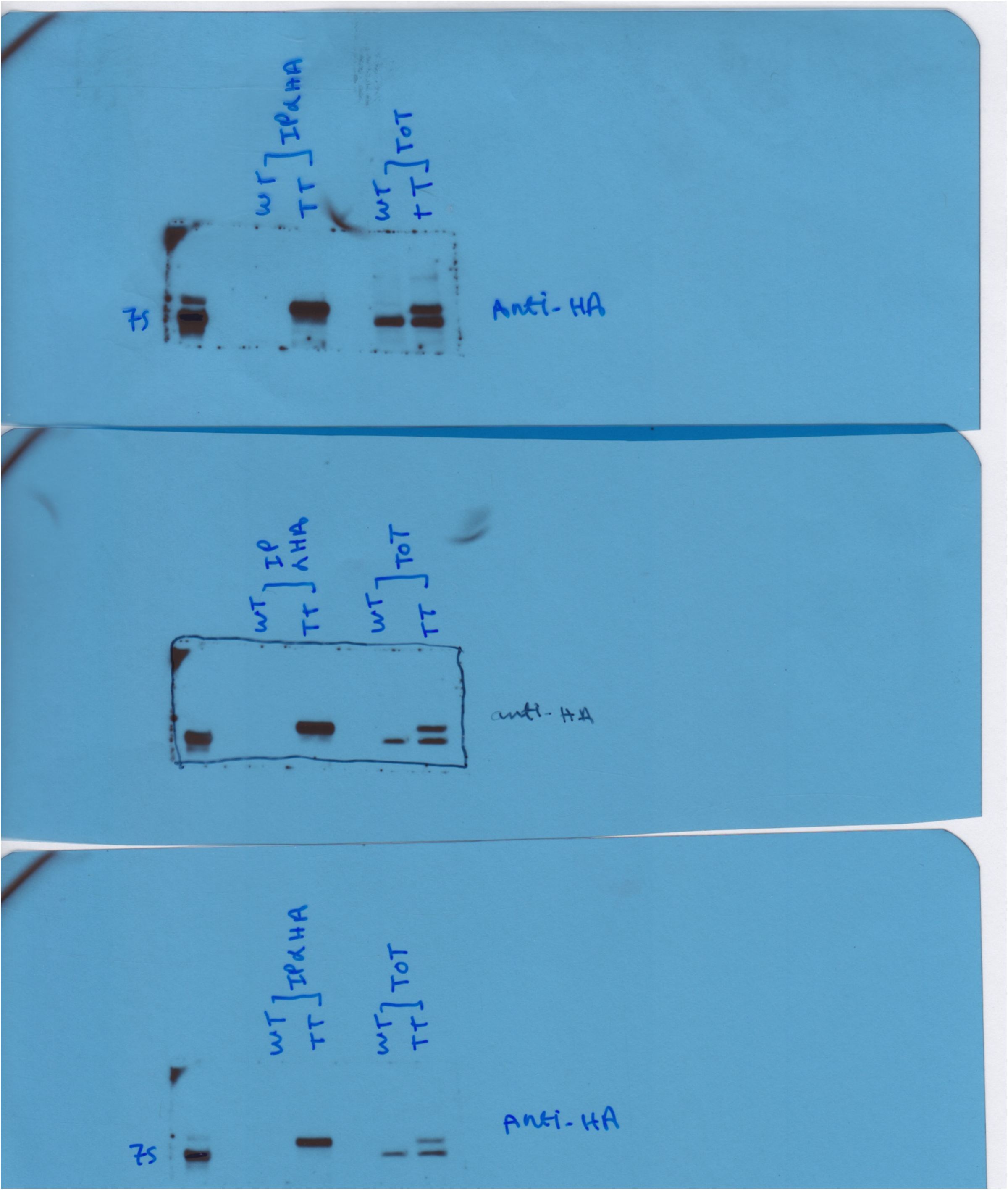
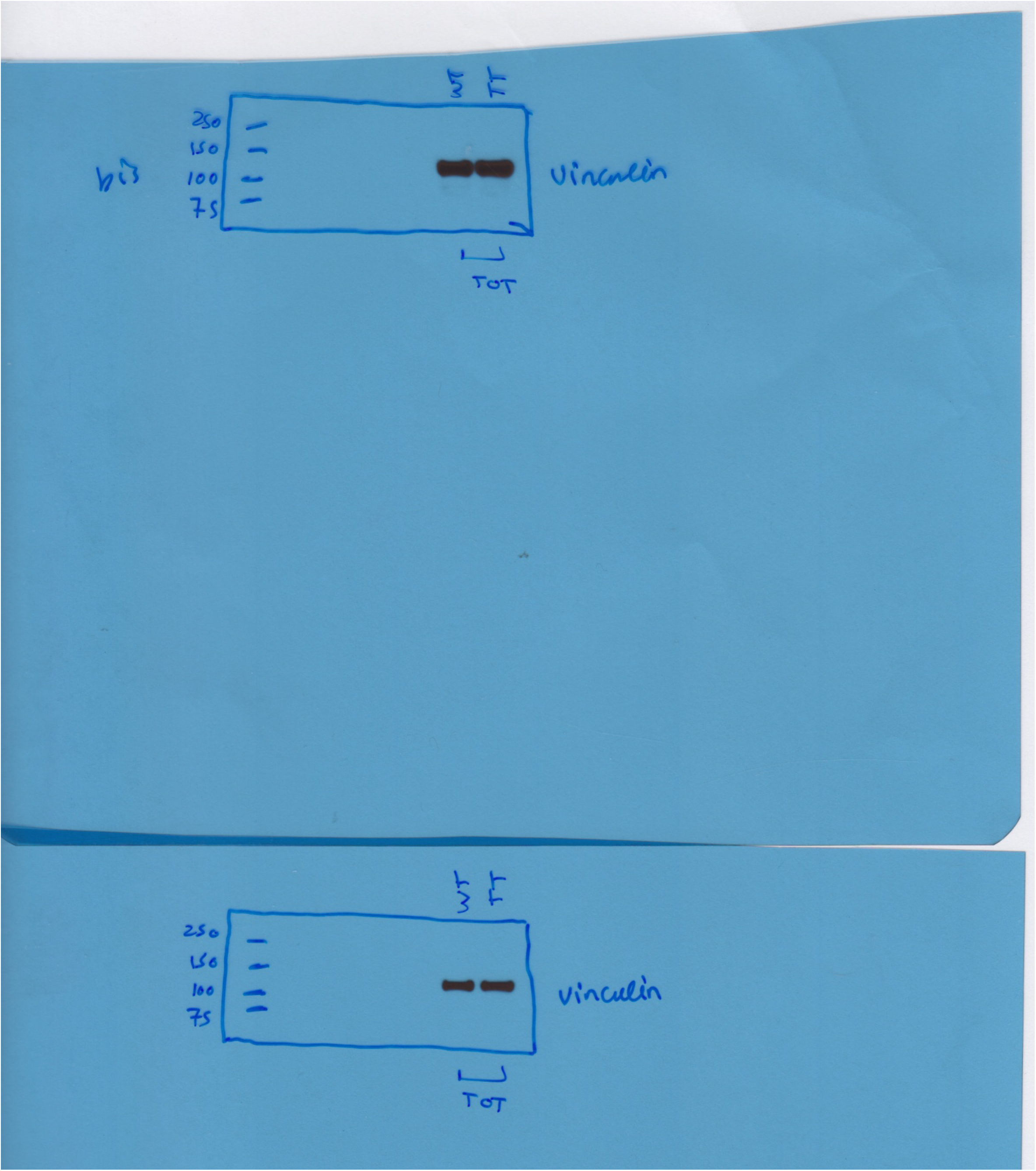
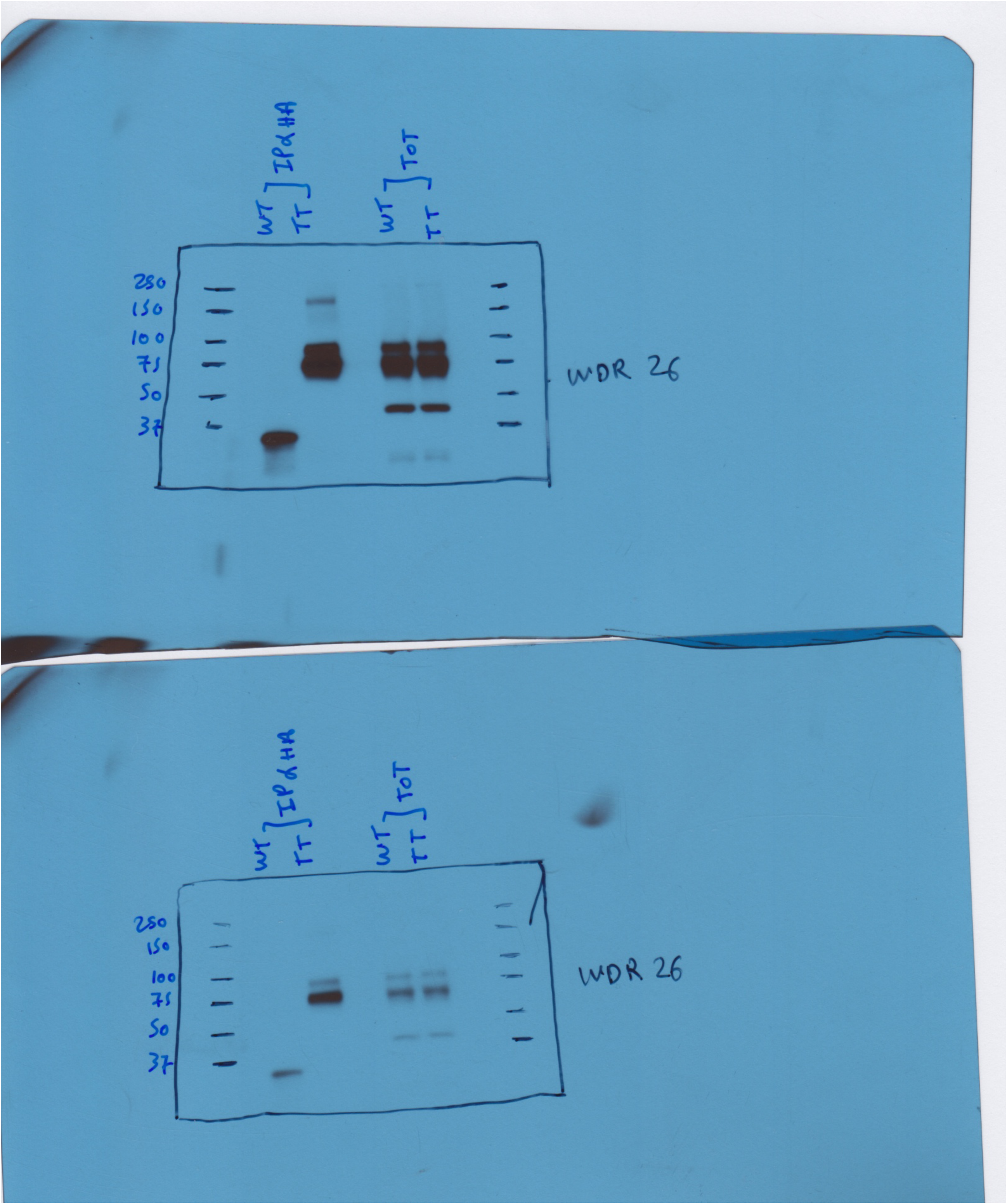
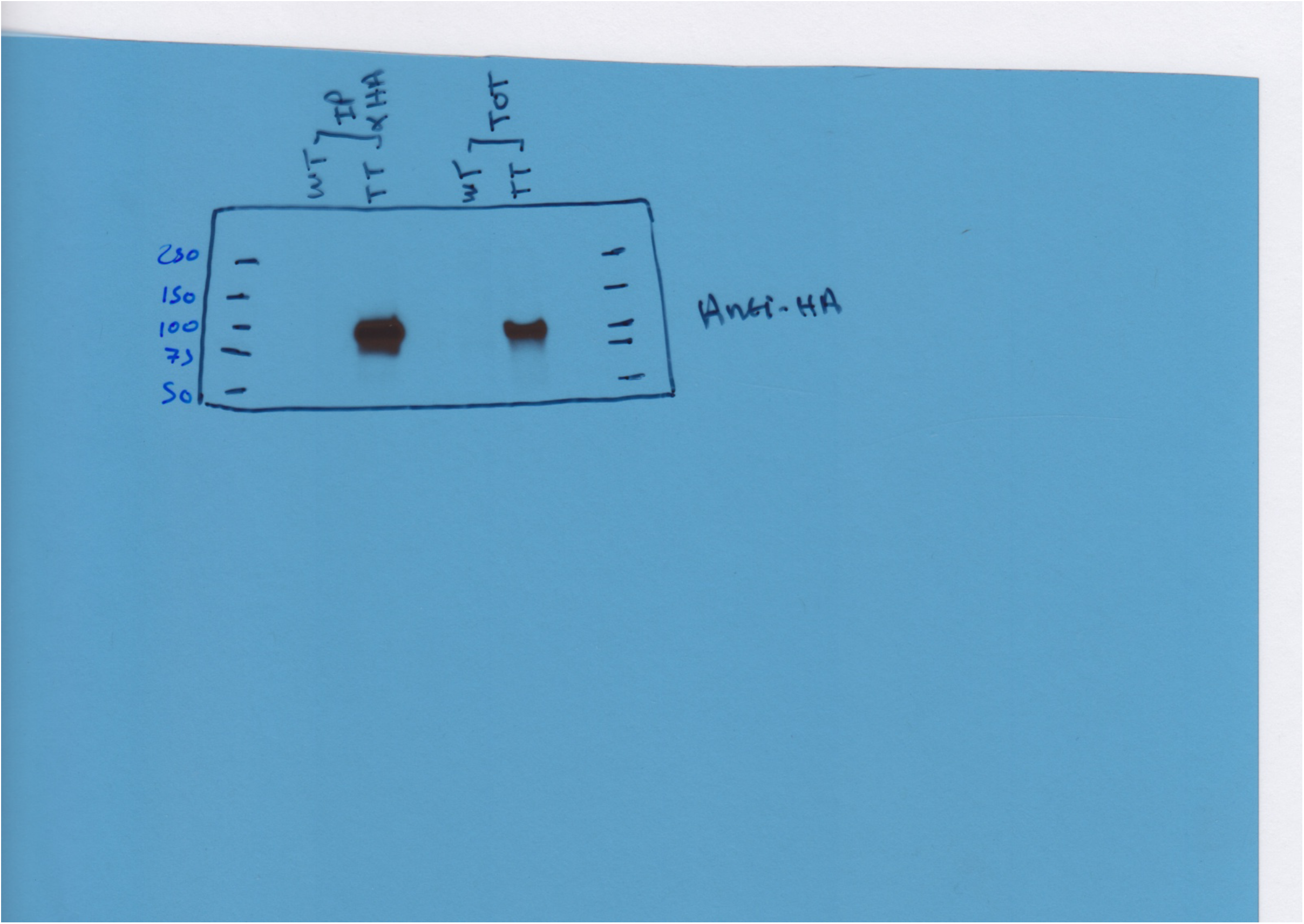
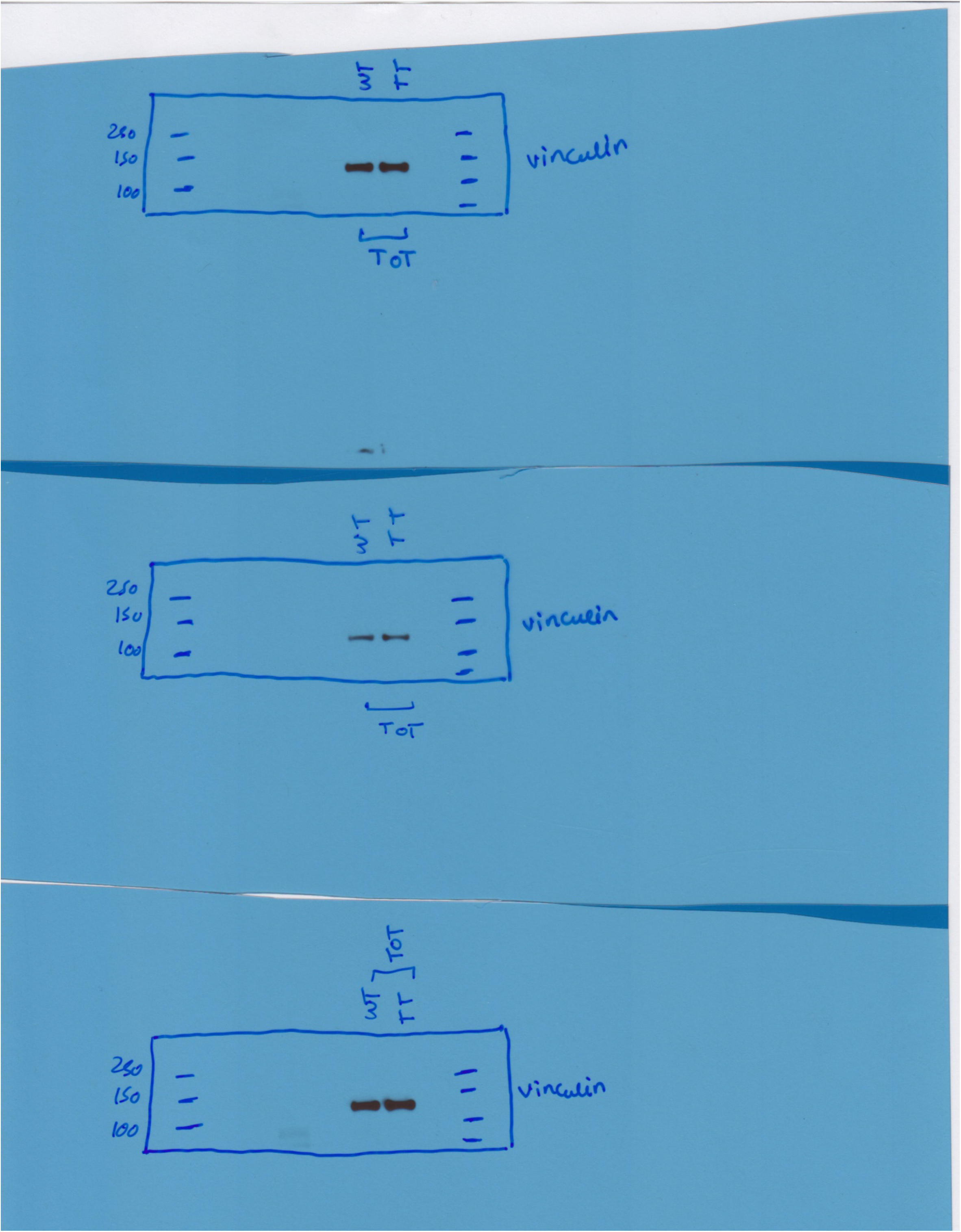
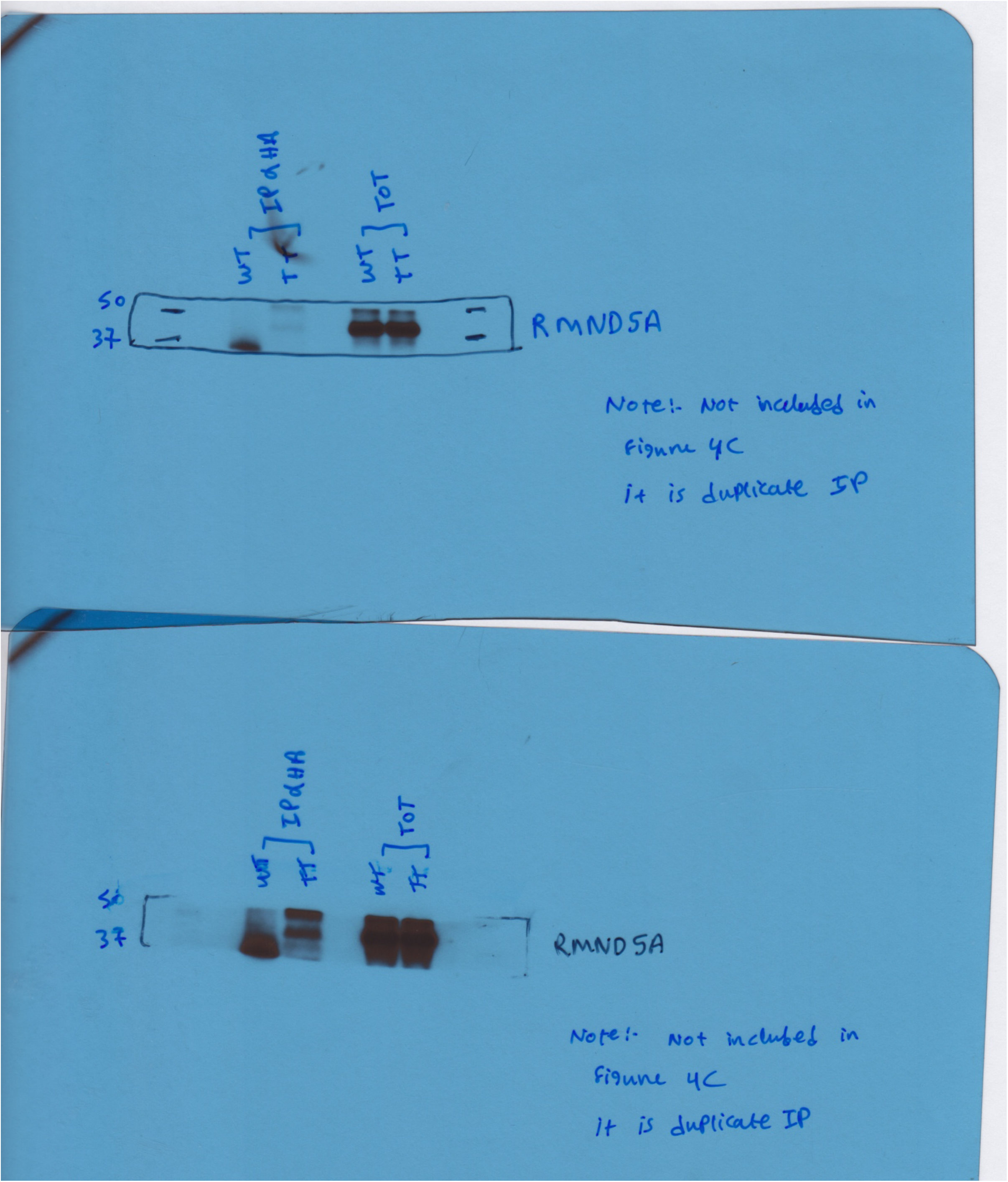
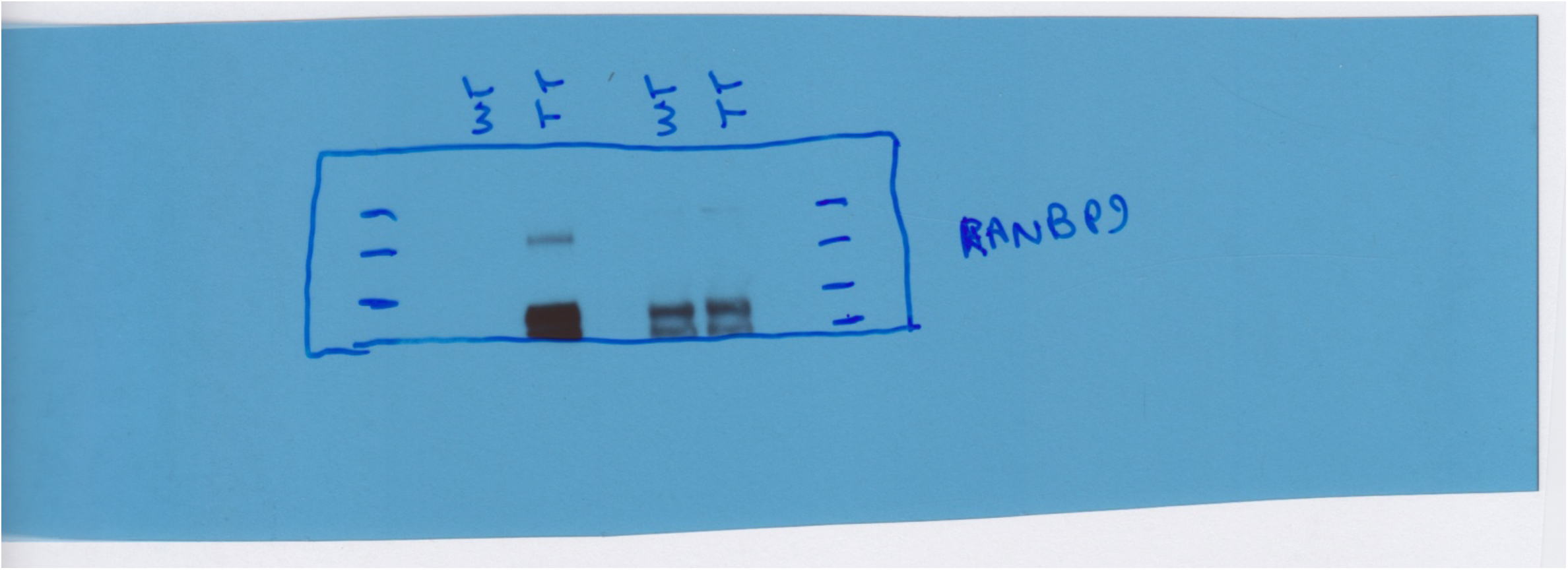
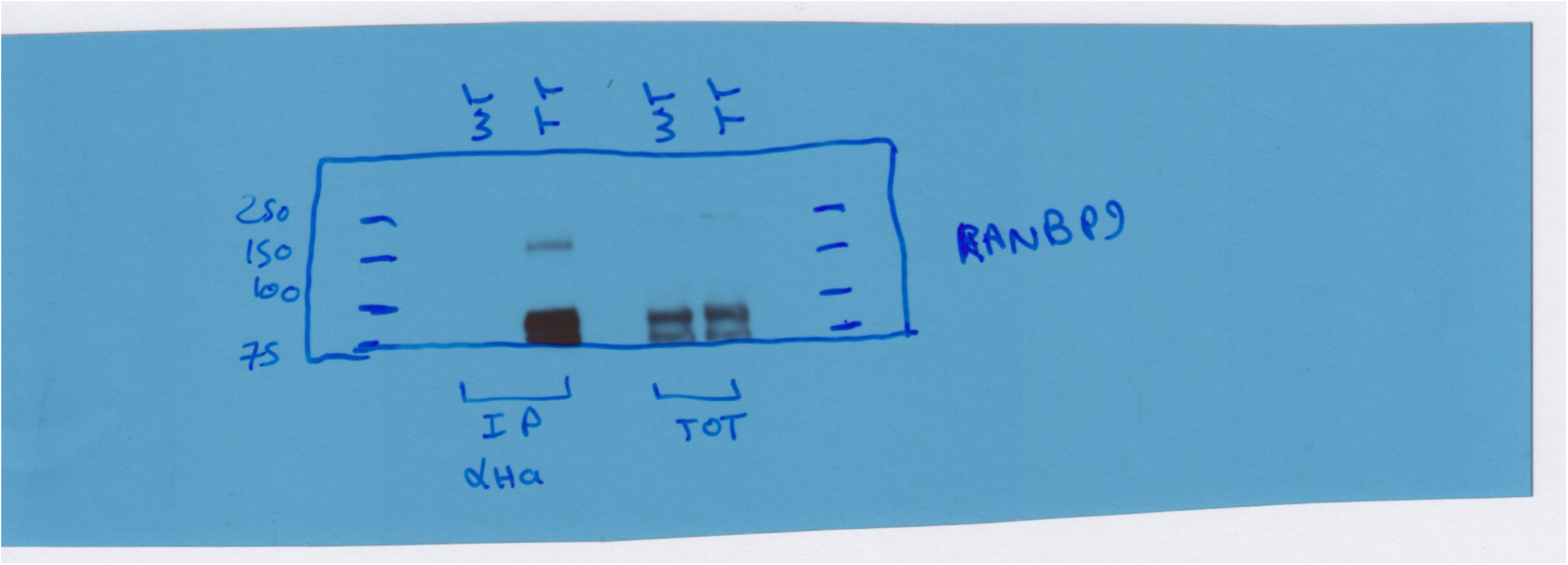
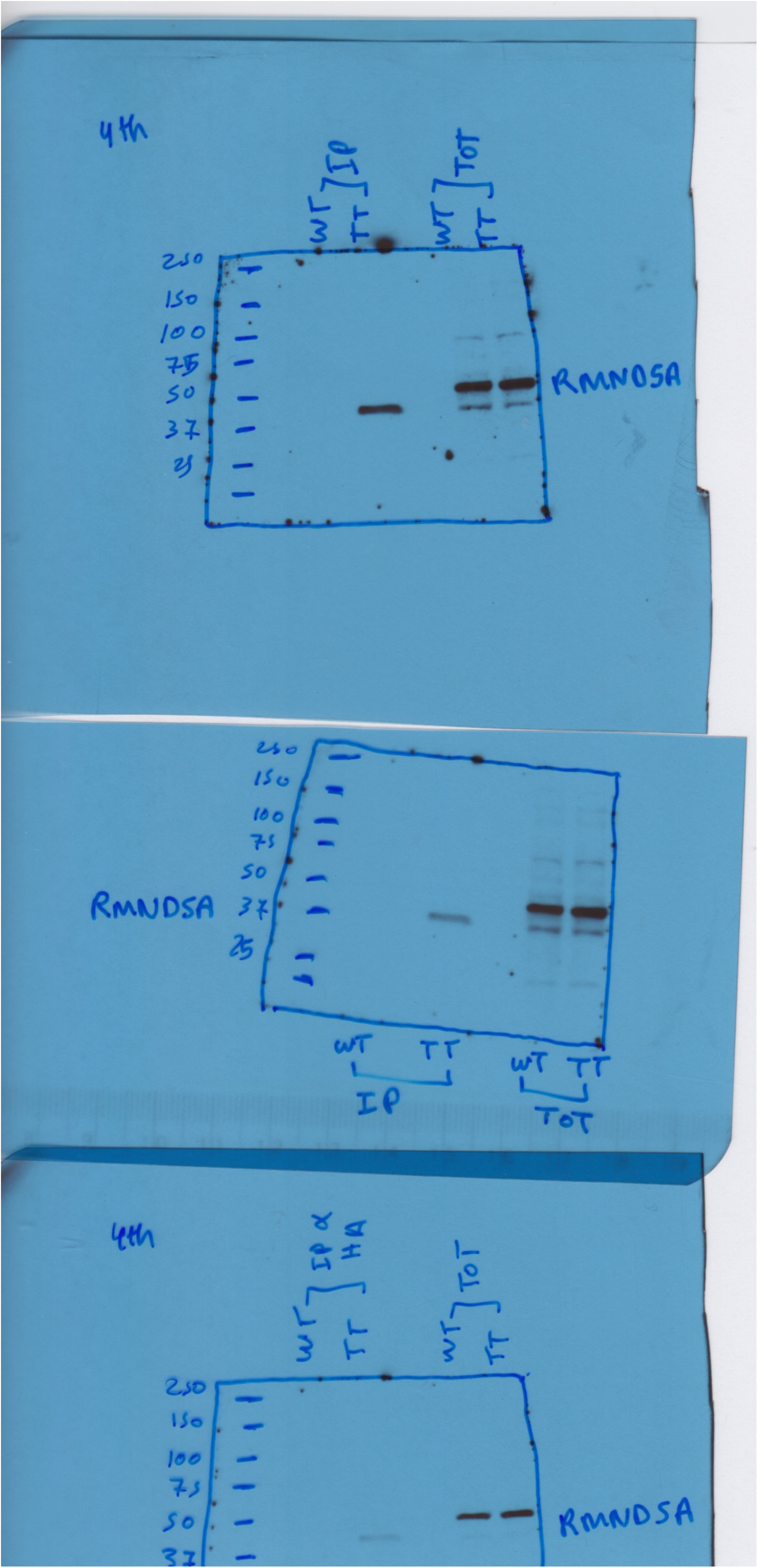
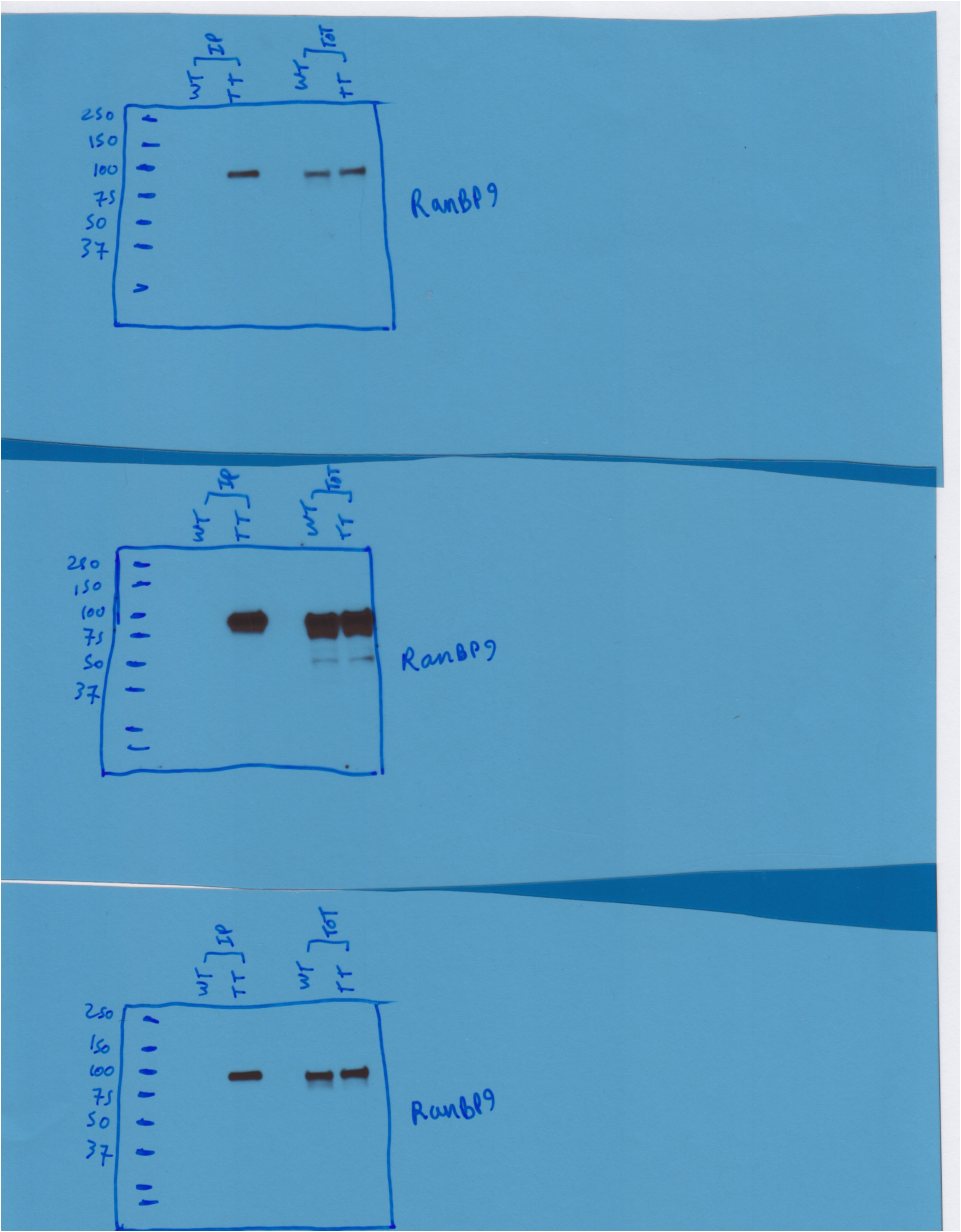
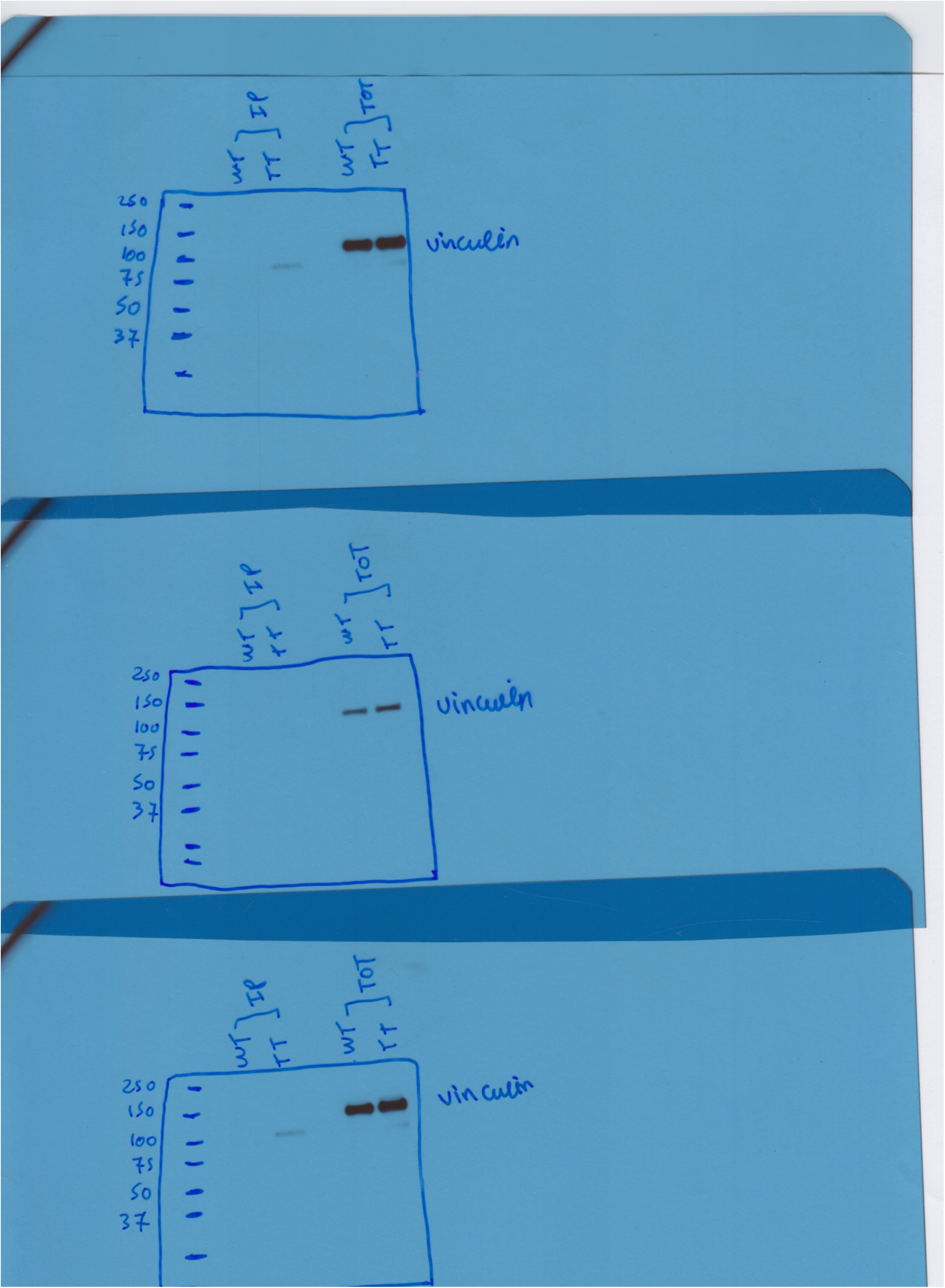
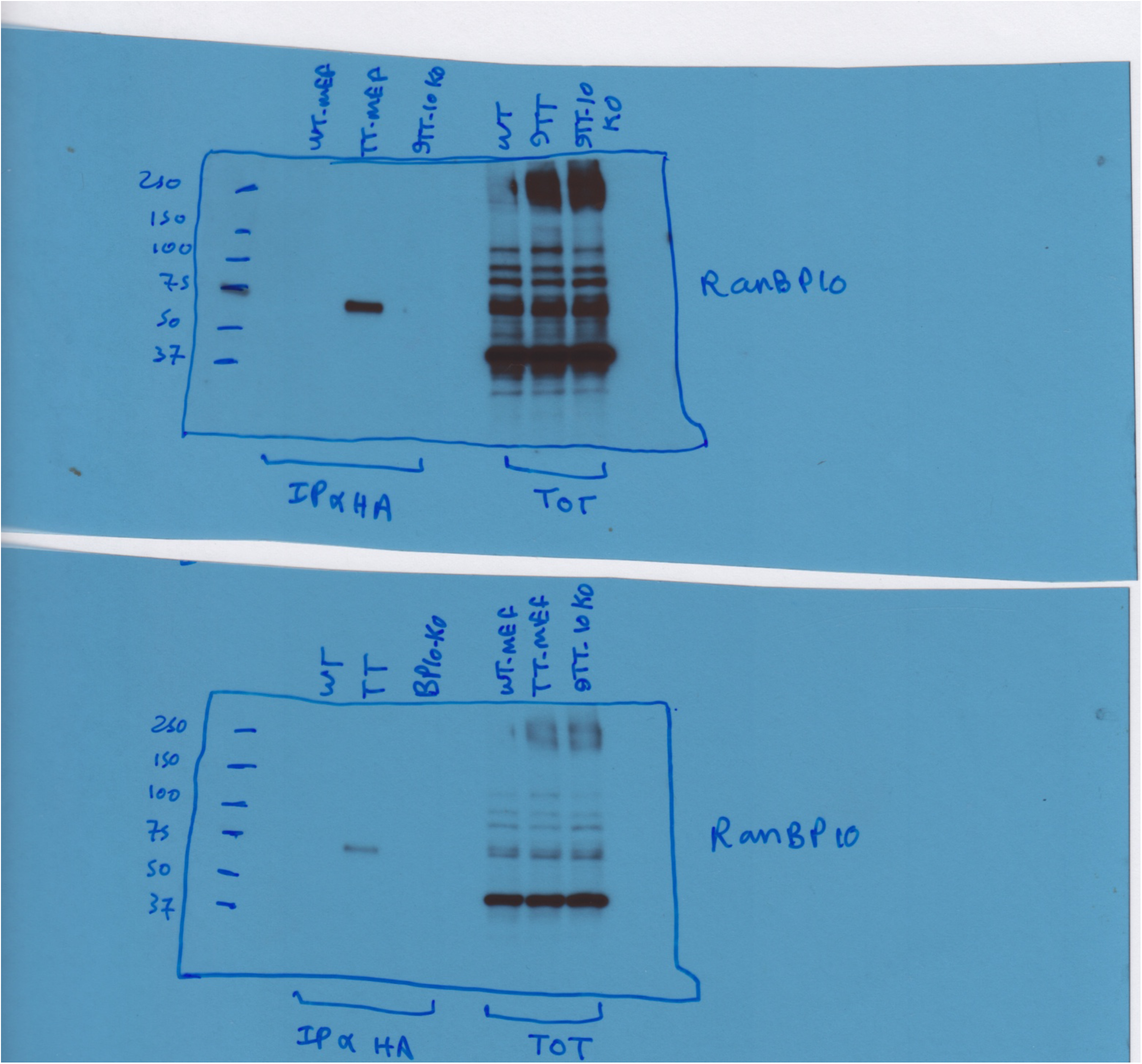
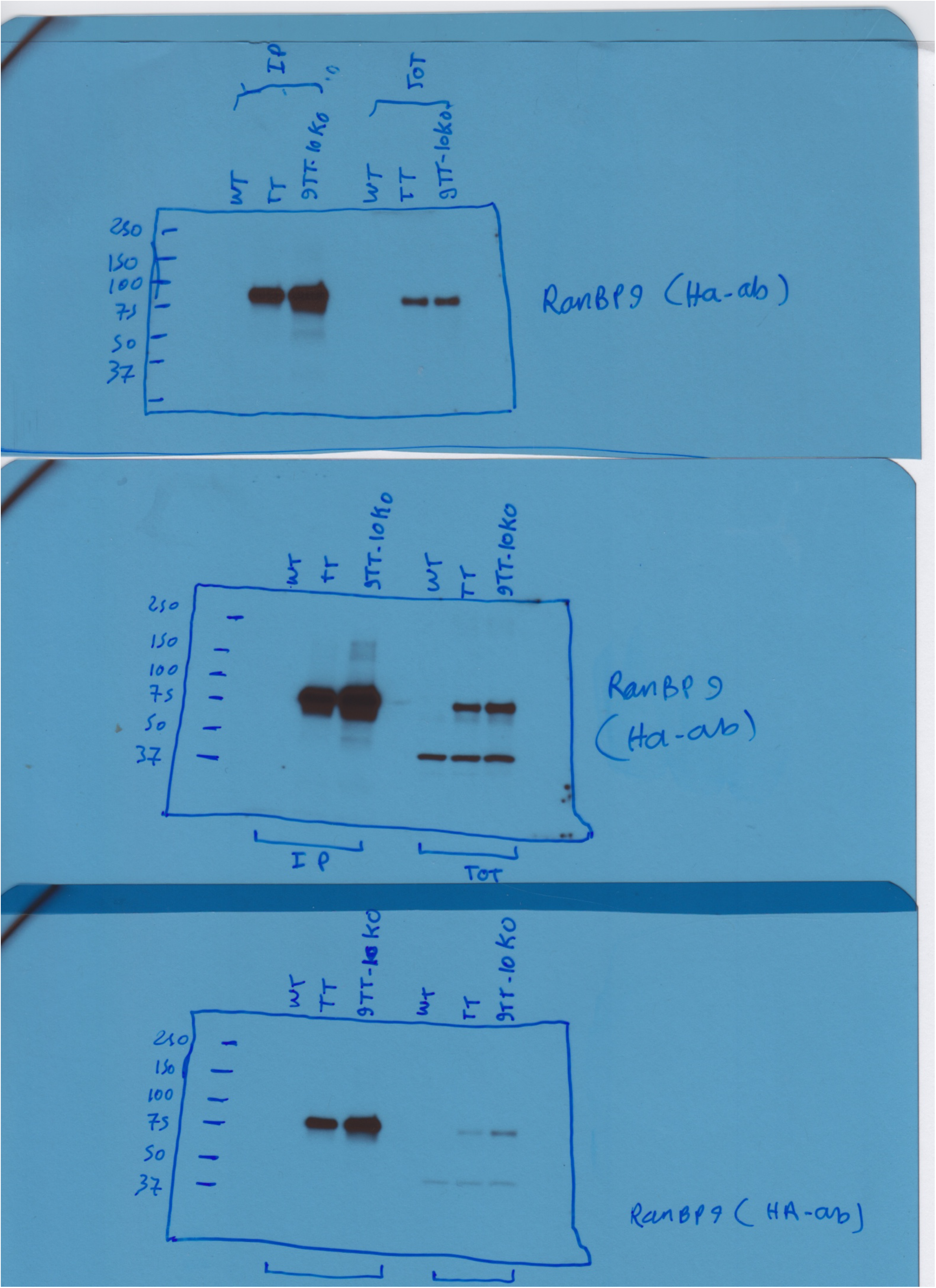
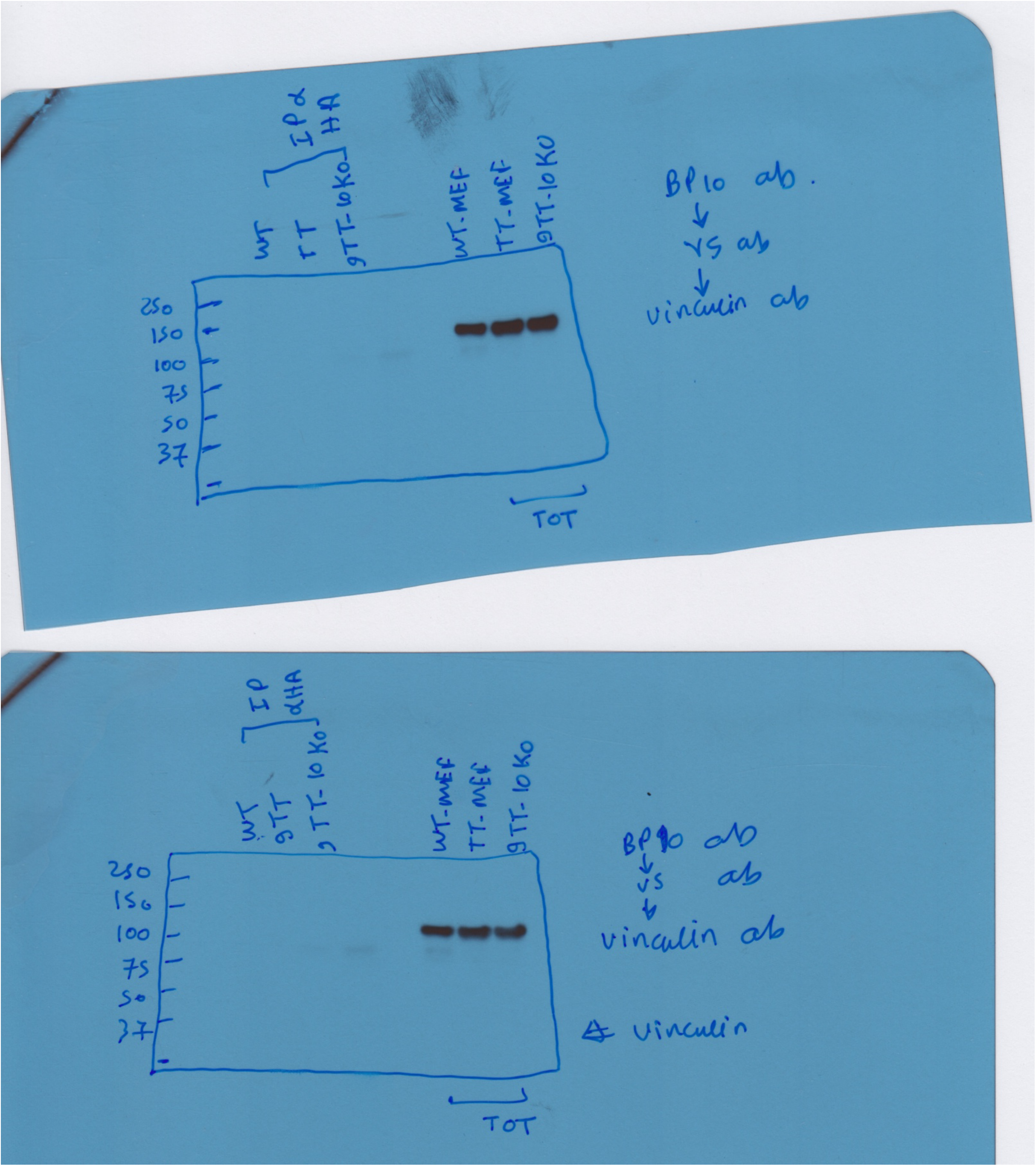
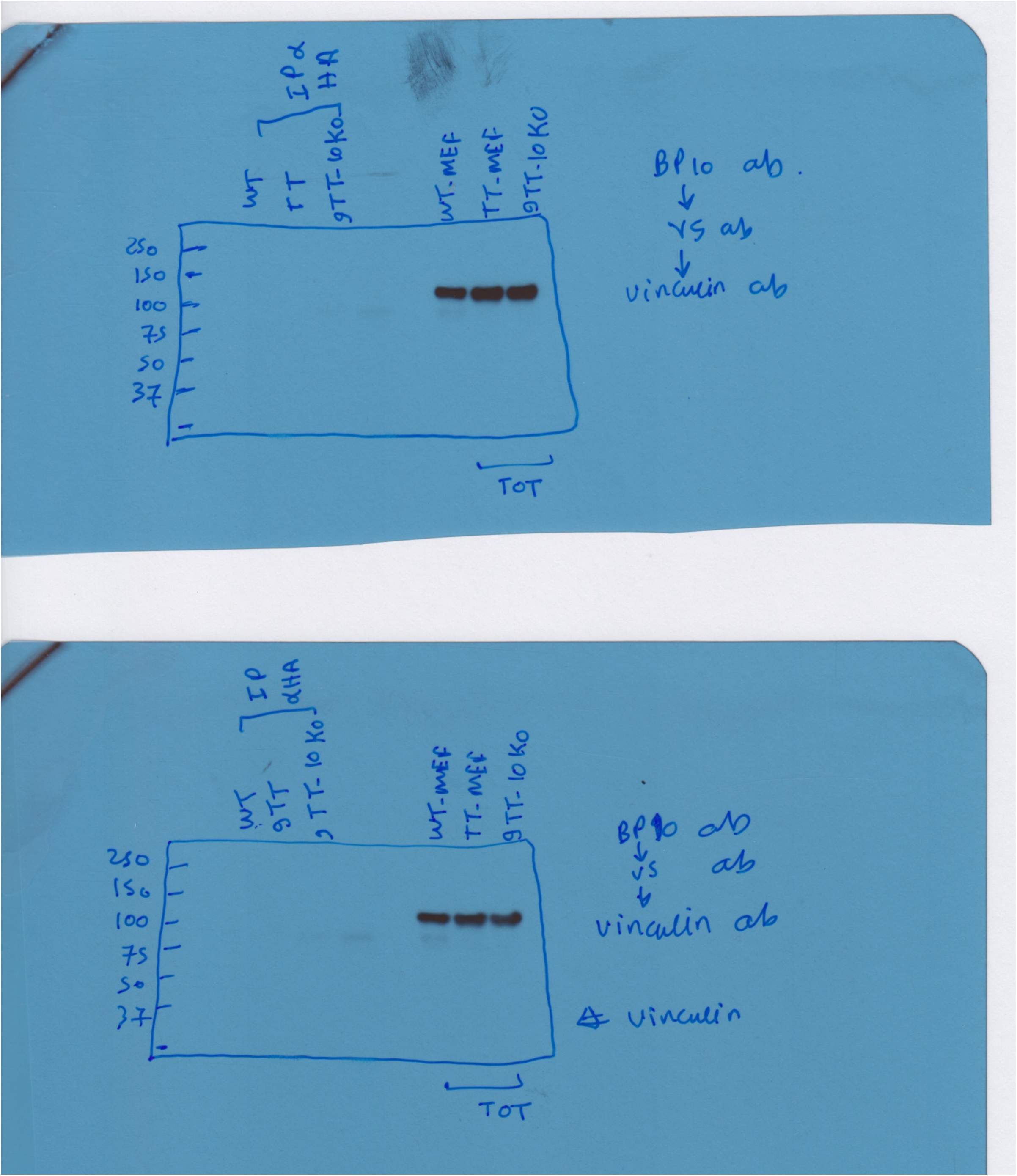
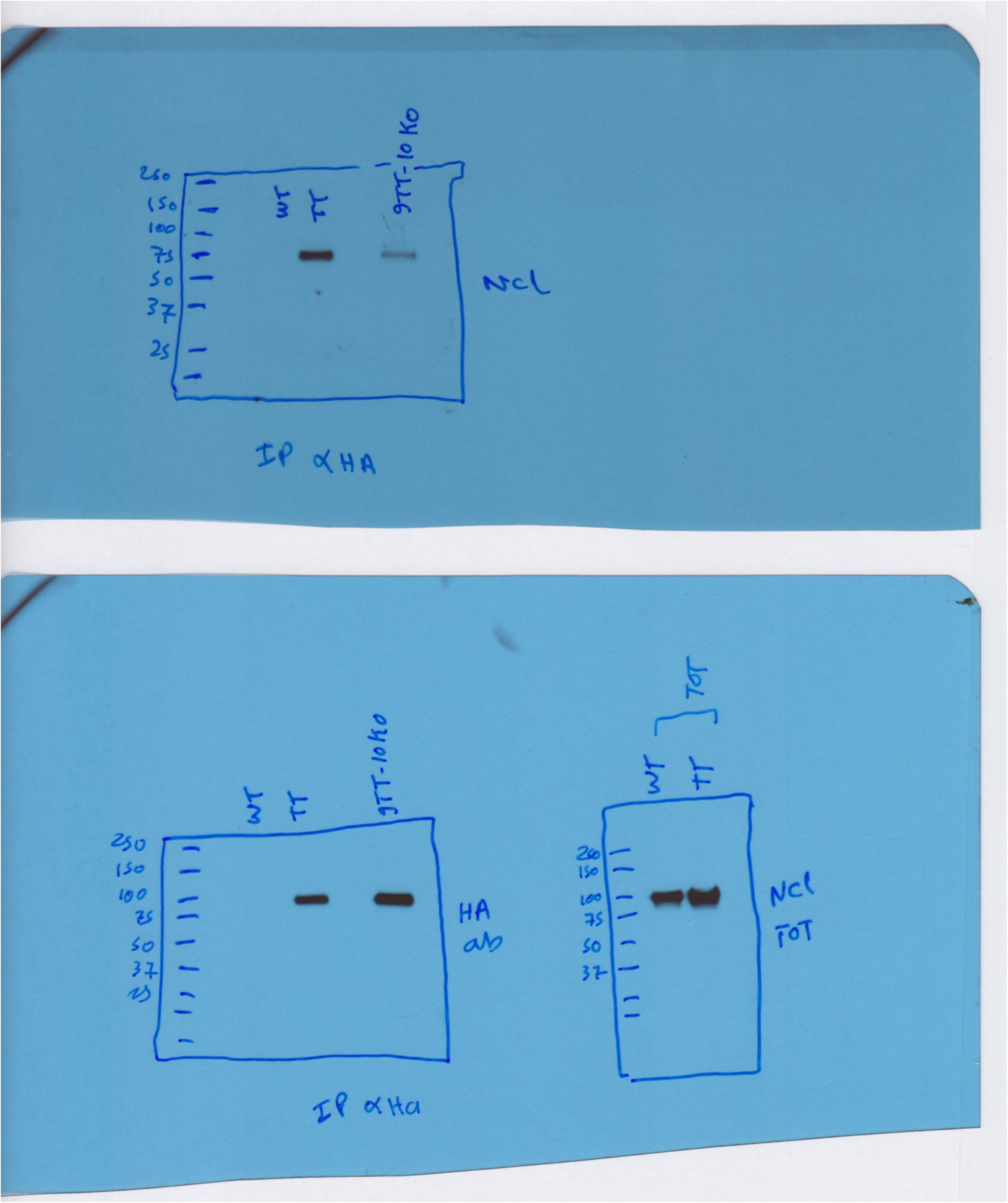
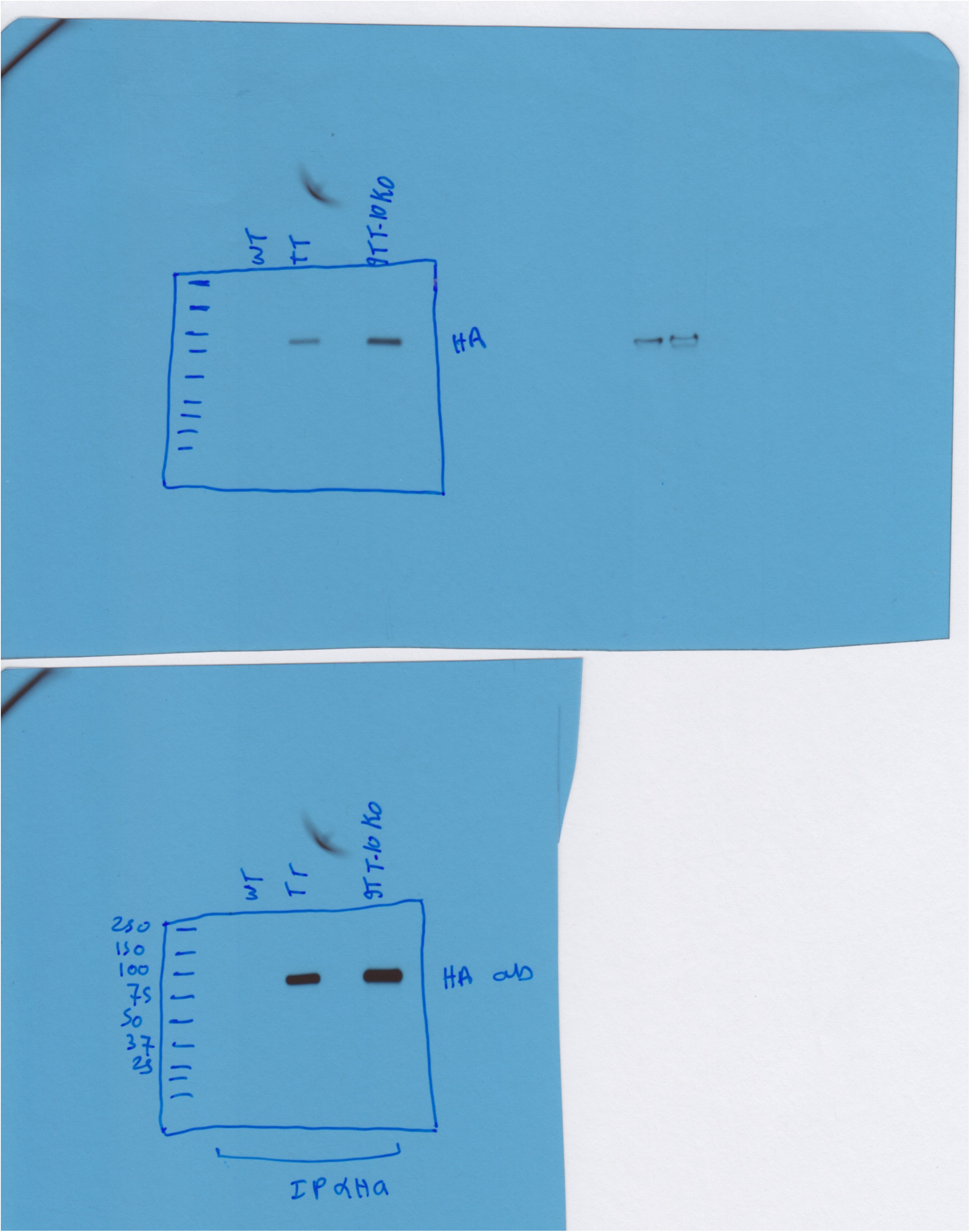
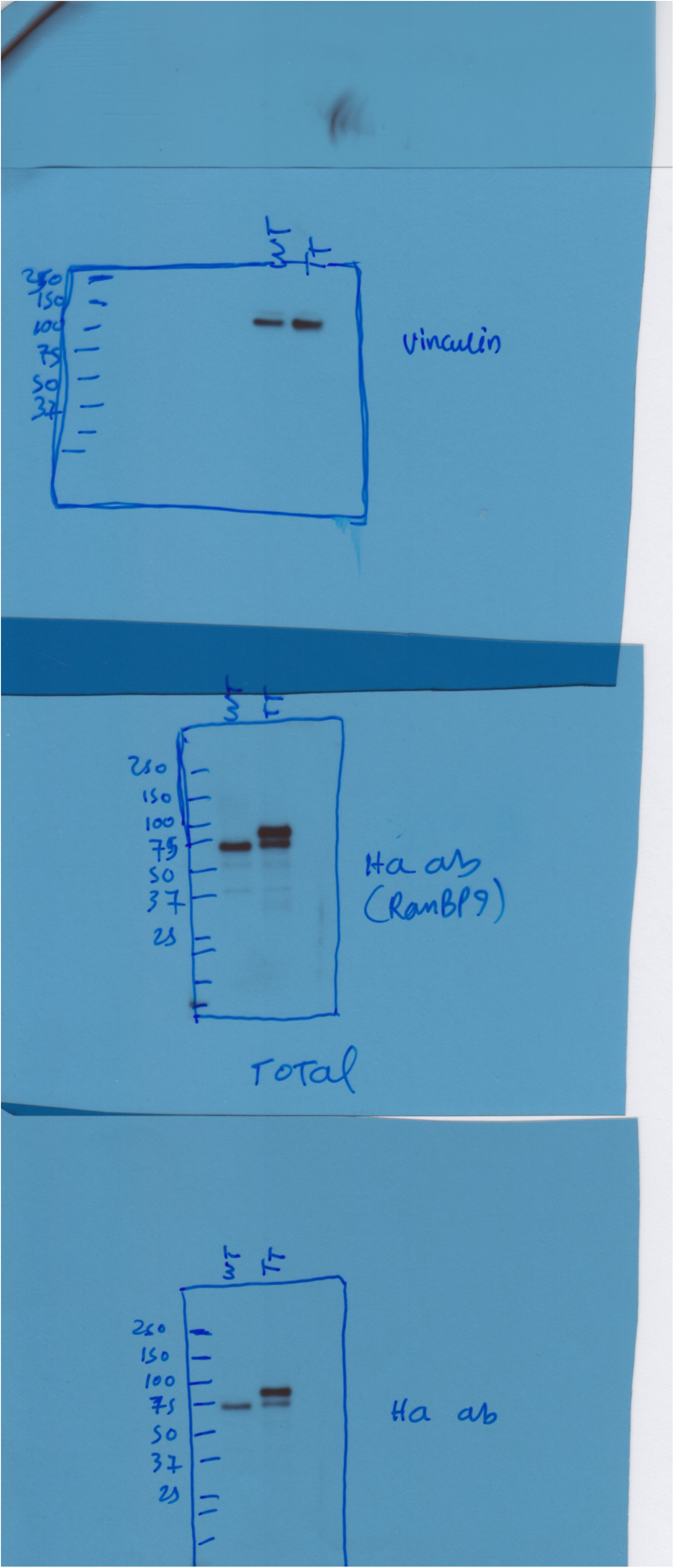

